# Memory encoding reprograms neuronal transcriptional responses via durable chromatin remodeling

**DOI:** 10.64898/2026.07.29.741555

**Authors:** Yuxi Ke, Xiaochen Sun, William J. Greenleaf, Mark J. Schnitzer

## Abstract

The mammalian brain’s long-term memory circuits integrate information from prior and new experiences. The medial prefrontal cortex (mPFC) has a crucial role in this process and can reliably store information for weeks to months in rodents and over years in humans. To maintain information over these extended timescales, the neural encoding of remote memories involves persistent synaptic, transcriptional, and epigenetic changes that outlast the more transient forms of molecular activation that occur in the initial minutes to hours of memory storage. However, whether these persistent effects include long-lasting changes to chromatin structure and whether chromatin states mainly reflect prior episodes of neural activation or retune transcriptional responses to future bouts of activation remain unknown. Here we show that mPFC neurons engaged during the initial formation of memory undergo progressive changes in chromatin accessibility over the first four weeks of memory storage, evincing long-lasting modifications to the genetic programs activated during subsequent memory retrieval. Our experiments involved genetic trapping and single-cell multiomic sequencing analyses of mouse mPFC engram neurons activated during contextual fear conditioning. In the absence of subsequent memory recall, memory storage-related changes to chromatin structure were modestly reflected in gene expression patterns at 7 and 28 days after fear conditioning. However, upon memory recall, the genetically trapped engram neurons executed distinct transcriptional programs from those of other neurons of the same genetic types, suggesting that chromatin rearrangements arising during remote memory storage alter the transcriptional control logic by which engram neurons respond to new experiences. These metaplastic changes to transcriptional programs preferentially affect gene-regulatory and post-transcriptional control mechanisms, show substantial enrichment for transcription factor motifs related to neural development and cell-state regulation, and downregulate the neuron’s transcriptional responses to future excitation. Thus, rather than merely preserving a molecular record of prior learning, chromatin architectural changes in engram neurons occur over timescales of weeks and appear, in part, to repurpose conserved regulatory machinery to dampen the extent to which these neurons will engage in further information storage. Based on these findings, we propose that chromatin structural changes provide a slow-timescale component of neural computation that reduces interference between the representations of different memories.

## Introduction

In the days to weeks after learning, long-term memories become increasingly dependent on circuits of the medial prefrontal cortex (mPFC), which can store information for months in rodents and over years in humans^1^. Within these circuits, engram cells, neurons activated during learning and reactivated during memory recall, are widely considered to provide a cellular substrate of long-term memory^2^. After contextual fear conditioning, mPFC engram cells are relatively silent or dispensable during memory recall episodes that occur soon after learning but become more active and are necessary for memory recall at remote time points long after the initial memory storage^3,4^. This delayed necessity for mPFC engram neurons poses a vexing biological problem: how do these neurons acquire durable states that are initially silent but later support remote memory retrieval? More broadly, how can memory circuits remain sufficiently plastic to store new information, yet sufficiently stable to preserve information across extended timescales?

Although stability is as fundamental to memory as plasticity, far more experimental attention has been given to how memories are initially written than to how they are maintained. Many molecular studies of memory have focused on activity-regulated transcription, including by immediate early genes, such as AP-1-driven programs that operate in the minutes to hours after a learning episode^5^. In contrast, the slower regulatory machinery that stabilizes cellular states over the subsequent days to weeks remains poorly understood. Developmental and lineage-defining transcriptional programs provide canonical examples of durable biological state regulation, but are often treated in the field as being separate from the transcriptional programs underlying memory.

Chromatin is a prime candidate for enabling the persistence of remote memories^6,7^. Learning can alter DNA methylation^8,9^, histone acetylation^6,10^, enhancer accessibility^11–13^, and 3D genome organization^14,15^, and perturbing these facets of chromatin can either disrupt or strengthen a long-term memory^9,16,17^. Yet, how chromatin changes functionally contribute to memory storage and retrieval remains unresolved. To date, chromatin’s role is largely framed as one that supports activity-regulated transcription, carrying forward the effects of neural spiking so that subsequent memory recall events can induce appropriate forms of gene expression^11,14,18–21^.

However, memory retrieval involves more than just a simple playback of neural activity that occurred during learning. When an animal recalls a memory, engram neurons are reactivated, and this reactivation can itself modify the memory^22–25^. Yet, this updating of previously stored information can destabilize it, so reactivated neurons must balance the need to respond to experience with the need to preserve existing memories^26,27^. This suggests that long-term memory storage may require mechanisms that regulate how neural activation is translated into plasticity-related gene expression, raising the possibility that chromatin may guide which genes are engaged upon neural reactivation and the levels of their expression^28^. Such an experience-dependent role for chromatin regulation might predict changes not only in plasticity effector genes that implement changes in synaptic function, excitability, or structure^4,29^, but also in upstream transcriptional^30^ and post-transcriptional^31–34^ regulators that control how effector programs are selected^35^, amplified, attenuated, or rerouted after reactivation^36,37^.

Based on this reasoning, we asked whether cortical engram neurons preserve chromatin changes after an initial memory encoding event or acquire chromatin states that alter transcription responses at memory recall. To approach this question, we applied activity-dependent genetic tagging of mouse engram neurons^38^ with joint single-nucleus RNA and ATAC multiome sequencing to profile mPFC neurons at 7 and 28 days after contextual fear learning, either with or without memory recall at these two time points after learning. This design allowed us to differentiate gene expression patterns in engram cells in response to memory recall events from those occurring without recall. We found that at 28 days after learning and in the absence of recall, excitatory mPFC engram neurons show modest transcriptional but large chromatin changes that were absent at the 7-day time point. After memory recall, engram neurons differed strongly from non-engram neurons in their transcriptional responses. This gene expression modulation engaged not only plasticity effectors but also upstream regulatory machinery, consistent with the idea that chromatin shapes gene responsiveness at recall, rather than simply relaying past neural activity.

At one week after learning, chromatin signatures in engram neurons showed residual AP-1-associated accessibility increases consistent with recent neural activation during learning. By four weeks, these AP-1-associated changes had receded and were replaced by chromatin accessibility changes that were enriched for transcription factor motifs linked to developmental and conserved cell-state regulatory programs. Memory-related chromatin information was distributed across both developmental and adult regulatory elements. Strikingly, at one week after learning, the chromatin state was not associated with gene expression patterns that were specific to engrams during subsequent memory recall, but at four weeks the chromatin state was predictive of such gene expression patterns.

Further, inhibitory neurons exhibited a distinct regulatory strategy from that of excitatory neurons. Inhibitory cells showed stronger persistent transcriptional differences in the absence of recall and weaker recall-dependent modulation of gene regulation. Nevertheless, inhibitory neurons shared broad chromatin and transcriptional engram signatures with those of excitatory neurons, including enrichment of developmental transcription factor programs at four weeks. Together, these results support a model in which cortical memory consolidation recruits a slowly maturing regulatory state that shapes how neurons transcriptionally respond to future recall.

Overall, we conclude that chromatin architectural changes in engram neurons emerge over timescales of weeks and use conserved regulatory machinery to reduce the amplitude with which engram neurons will be activated by future events. Notably, classic models of memory storage, such as the Hopfield model, recognize the perils of ‘memory interference’, in which the storage of new memories can jeopardize the persistence of older memories. Based on our results, we propose that chromatin structural changes serve in part to reduce interference between the representations of different memories, by dampening the engagement of engram neurons to incoming information that promotes new memory storage.

## Results

### Multiomic profiling of mPFC engram neurons at one and four weeks after fear learning

We set out to track transcriptional and chromatin changes in mouse mPFC engram neurons across several weeks during which systems consolidation occurs following fear conditioning^39,40^. To this end, we used drug-inducible, activity-dependent genetic tagging to label the nuclei of engram neurons activated during contextual fear conditioning with a nuclear-envelope-targeted form of the green fluorescent protein (GFP) (**Methods**; **Fig. 1a**). We isolated the nuclei of mPFC neurons at either 7 d or 28 d after fear conditioning, either without or 60 min after a memory recall session in which the mice revisited the conditioning context immediately prior to tissue collection. We then sorted the nuclei into GFP+ and GFP– fractions, respectively representing engram and non-engram mPFC neurons (**Supplementary Fig. 1a**). Importantly, the mice exhibited conditioned freezing levels that were comparable right after fear conditioning and in the memory recall sessions at 7 and 28 days (**Fig. 1b**).

**Figure 1.**
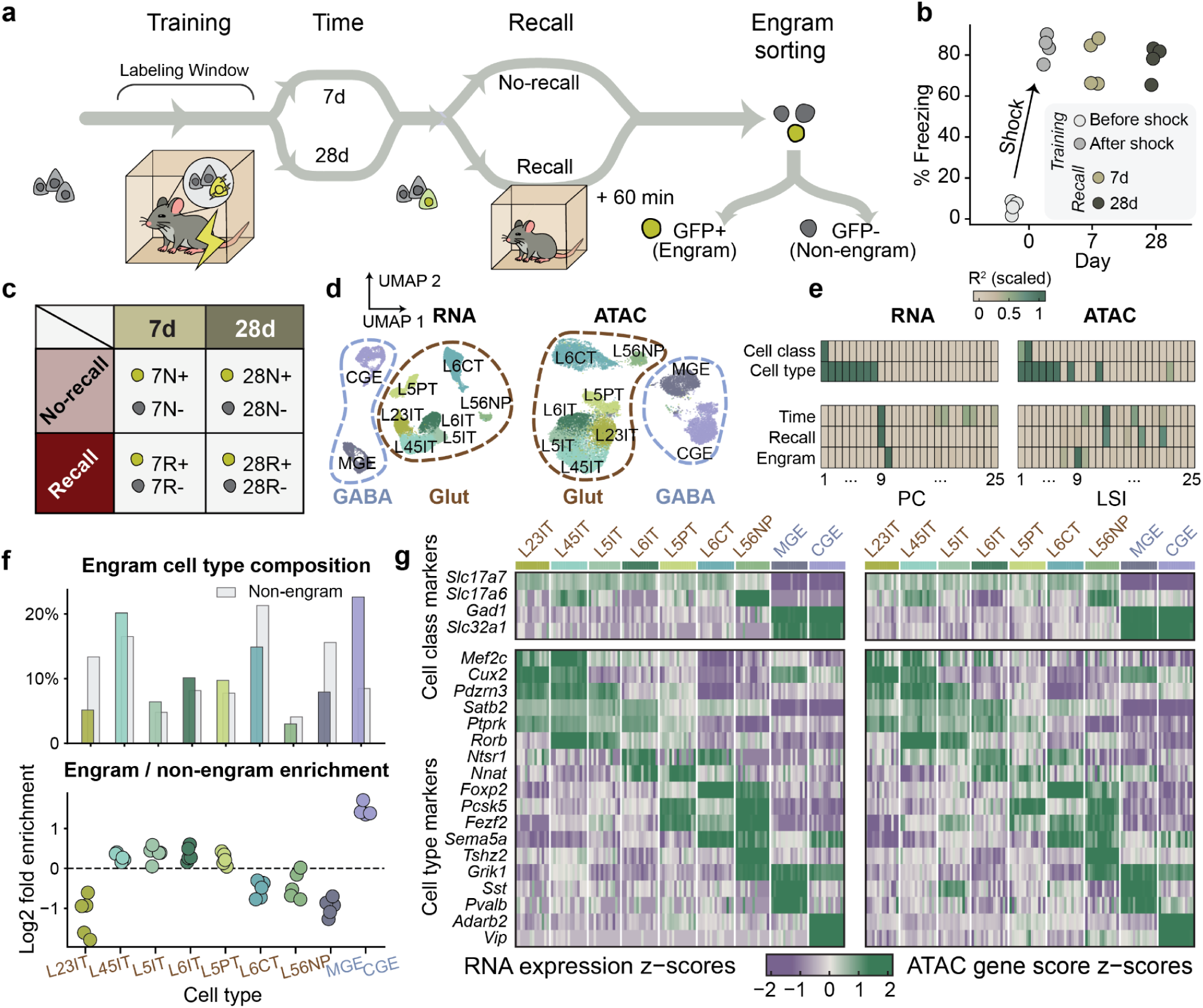
Multiomic profiling of mPFC engram neurons at 7 and 28 days after fear memory encoding reveals cell-type information and enrichment of engram cells in specific cell types. **a)** Schematic of the experimental workflow. Using a Fos-TRAP2; Sun1-GFP mouse line, mPFC neurons activated during contextual fear conditioning were genetically trapped and received a nuclear fluorescent tag. At either 7 or 28 days later, mice were either re-exposed to the conditioning context (Recall group) or kept in their home cage (No Recall group). 60 min later, we collected the cell nuclei of mPFC neurons and sorted them into GFP+ (engram) and GFP– (non-engram) populations. **b)** When returned to the conditioning context at either 7 or 28 days after the initial conditioning session, mice exhibited robust freezing responses, confirming the retention of stable fear memories across the time points used for multiomic profiling. X-axis, training: before and during shock delivery; recall: 7 and 28 days after training. Y-axis, percentage of time mice spent freezing in a five-minute window. Each datum shows the results from an individual mouse. **c)** Table showing the experiment’s factorial design across time (7 *vs.* 28 days), memory recall condition (Recall *vs.* No recall), and engram status (Engram *vs.* Non-engram cell), yielding a total of 8 conditions. Most biological replicates consisted of pooled tissue from 4 mice, except for one replicate pair in the 7-day recall condition (7R+/−), which consisted of pools of 3 mice (**Supp. Table 1**). Each mouse pool generated a matched pair of engram and non-engram samples, except for one replicate pair in the 28-day no-recall condition (28N+/−), where engram and non-engram samples originated from two independent mouse pools. **d)** UMAP plots of RNA (*left*) and ATAC (*right*) profiles across all nuclei show that differences associated with neuron-type dominate the global space of variations across cells. The plots show clear separations between glutamatergic and GABAergic neuron classes, as well as finer divisions into established cortical cell types. Abbreviations: L23IT, layer 2/3 intratelencephalic neurons; L45IT, layer 4/5 intratelencephalic neurons; L5IT, layer 5 intratelencephalic neurons; L6IT, layer 6 intratelencephalic neurons; L6CT, layer 6 corticothalamic neurons; L5PT, layer 5 pyramidal-tract neurons; L56NP, L5/6 near-projecting glutamatergic neurons; CGE, caudal-ganglionic-eminence–derived interneurons (comprising Vip, Sncg, Lamp5 subtypes); MGE, medial-ganglionic-eminence–derived interneurons (comprising Pvalb, Sst, Sst/Chodl subtypes). Results from each individual cell nucleus are represented by a single data point. **e)** The major dimensions of variability in the RNA and ATAC datasets capture cell type information, whereas subsequent dimensions capture information about cell state. Plotted are *R*^2^ values between each of the top 25 principal components of the RNA (*left*) or the top 25 LSI dimensions of the ATAC data (*right*) and either the cell type categories (*top plots*) or the time, recall, and engram conditions of panel **c** (*bottom plots*). Cell class: glutamatergic *vs.* GABAergic. Cell type: all 9 cell types in panel **d**. *R*^2^ values are scaled between 0–1 for all 40 dimensions used for analysis within each row. **f)** The set of engram cells spans all cortical neuron types but shows biased recruitment toward specific cell types. *Top*, Percentages of each cell type among the engram (GFP+; colored) and non-engram (GFP–; gray) nuclei sets. *Bottom*, Ratio of engram to non-engram cells within each cell type, plotted with a Log₂ scale on the *y*-axis. Horizontal dashed line: equal numbers of engram and non-engram cells. Each datum shows results from a single cell-sorting experiment. CGE-derived GABAergic neurons are the most enriched among the set of engram cells. **g)** Cell-type identity information is consistent across the RNA expression and ATAC datasets. The heatmaps show RNA expression (*left*) and ATAC (*right*) gene scores, z-scored across rows, for canonical marker genes across cell-type pseudobulks. The strong concordance in the results from the two modalities validated the cell-type assignments and enabled integrated analyses. Rows (marker genes); the “cell class” block includes glutamatergic and GABAergic neural markers, whereas the “cell type” block shows markers of finer cell-type divisions. Columns (biological pseudobulks) are ordered by cell types and grouped by sample.

For our data analyses, we designated the different sets of nuclei with a triplicate of variables to identify, respectively, the day on which the nuclei were isolated (either 7 or 28 days post-conditioning), whether the mice had undergone memory recall or not (N, R), and whether or not the nuclei expressed GFP (+, –). For example, 7R+ denotes GFP-expressing nuclei that were extracted at 7 days post-training from mice that underwent memory recall (**Fig. 1c**). In general, we collected two biological replicates per condition, with each replicate comprising a pool of four mice, with each pool generating a paired set of engram and non-engram samples (**Supplementary Fig. 1b; Supp. Table 1**).

Using the 10x Chromium platform, we generated multiomic sequencing data for a total of 17,669 nuclei after quality control and cell-type filtering (**Supplementary Fig. 1c-h**), yielding joint RNA and ATAC profiles that resolved major glutamatergic and GABAergic cell types (**Fig. 1d**). Glutamatergic classes included layer-specific intratelencephalic (IT) (L23IT, L45IT, L5IT, L6IT), corticothalamic (L6CT), pyramidal-tract (L5PT), and near-projecting (L56NP) neurons. GABAergic cell types included medial-ganglionic-eminence (MGE) and caudal-ganglionic-eminence (CGE) interneurons. These cell-type variations dominated the first 8 principal components (PCs) of either the RNA expression or the ATAC latent semantic indexing (LSI) data; cell state variations encoding the experimental condition were in the 9th reduced dimension **(Fig. 1e, Supplementary Fig. 1i,j**). All major cell types were present in the engram population, which was enriched for the GABAergic CGE cell type (**Fig. 1f**).

We generated biological pseudobulks by aggregating reads for nuclei coming from the same sample and cell type, yielding a total of 131 pseudobulks with at least 15 nuclei (**Supplementary Fig. 1k-n**). Marker genes and their chromatin accessibility scores confirmed the cell type assignments (**Fig. 1g, Supplementary Fig. 1h**). As expected, Fos RNA expression was higher in the GFP+ than in the GFP– population at 28 d with recall, consistent with engram cells having a higher probability of reactivation during remote memory recall^3^ (**Supplementary Fig. 1o**). For known cell-type markers, RNA expression levels and chromatin accessibility gene scores were highly correlated to each other across the pseudobulks, whereas these correlations were weaker for housekeeping genes (**Supplementary Fig. 2a-d**). Together, these results confirmed that our multiomic data were of high quality and consistent with canonical cortical neuron markers.

### Excitatory cortical engram neurons have distinctive responses at memory recall

Activity-regulated transcriptional programs unfold within a dominant transcriptional background that supports cell-type identity. To isolate subtle cell-state signals from this structure, we first focused on glutamatergic neurons, analytically modeled cortical layer-specific effects, and decomposed pan-glutamatergic transcriptional changes into a hierarchy of model-defined regulatory quantities (**Fig. 2a,b**). Within the high-dimensional spaces of RNA expression and chromatin accessibility peaks, we defined four regulatory ‘signals’ based on the gene expression or chromatin differences found between nuclei in different conditions.

**Figure 2.**
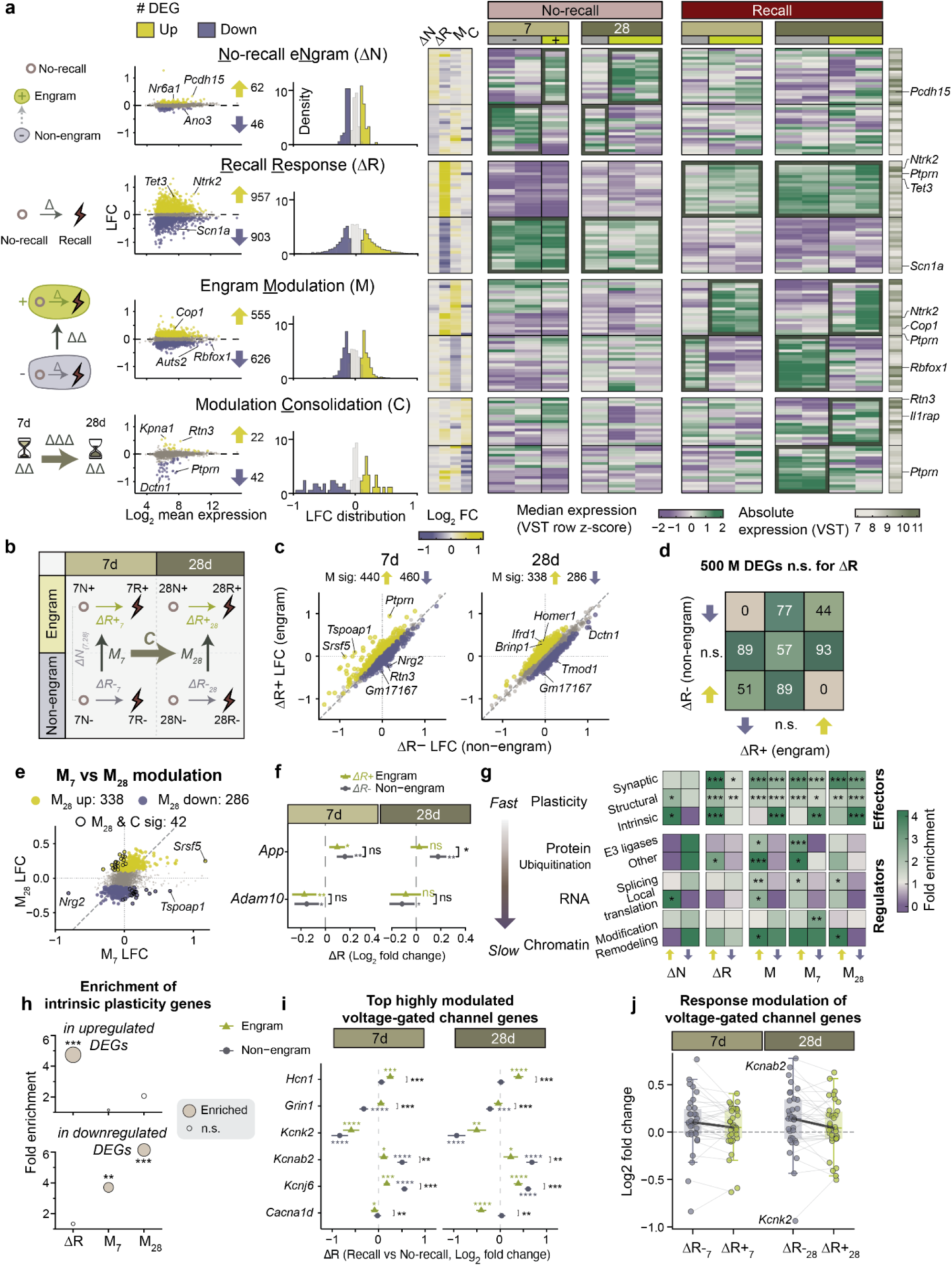
Glutamatergic engram cells primarily differ in how they respond to recall. **a)** Operational decomposition of transcriptional changes in glutamatergic neurons. Differential expression analysis across <u>n</u>o-recall engram (*ΔN*; engram vs non-engram without recall), recall response (*ΔR*; recall vs no recall), engram <u>m</u>odulation (*M*; *ΔΔR*, difference in *ΔR* between engram and non-engram neurons), and modulation <u>c</u>onsolidation (*C*; *ΔΔΔR*, temporal shift in *M* from 7 to 28 days). Yellow, significantly upregulated; purple, downregulated (s-value < 0.05, |LFC| > 0.1). Counts of differentially regulated genes (DEGs) are indicated for each quantity and direction by the up and down arrows. Left to right: MA plots show log₂ fold change (x-axis) versus mean expression (y-axis) with biologically representative genes labeled; histograms show density distribution (y-axis) of log₂ fold change (x-axis); heatmaps show log₂ fold change of the four quantities, median RNA expression (row z-score), and absolute expression level (VST, variance stabilized transformation from DESeq2, narrow bar on right) across all conditions. Row blocks, DEGs for each of the four quantities; rows, DEGs sorted by a score of |LFC| * -log10(s-value). Columns correspond to biological replicates (pools of four mice) of glutamatergic neurons, annotated by Recall, Time, and Engram status. Dark green boxes indicate expected expression patterns for each quantity. **b)** Conceptual summary of the NRMC framework. Schematic illustrating how <u>n</u>o-recall e<u>n</u>gram (*ΔN*), <u>R</u>ecall <u>R</u>esponse (*ΔR*), engram <u>M</u>odulation (*M*), and modulation <u>C</u>onsolidation (*C*) relate across experimental conditions. Light red circle, no-recall; dark red lightning bolt, recall. **c)** Engram neurons selectively reshape, rather than uniformly amplify, recall-evoked transcriptional responses. Scatter plots compare recall-evoked RNA log_2_ fold changes in engram (*ΔR+* LFC, y-axis) and non-engram (*ΔR-* LFC, x-axis) glutamatergic neurons at 7 d and 28 d (facet) after learning. The dashed diagonal marks equal recall responses in engram and non-engram neurons, so deviations from this line indicate engram-dependent modulation of the recall response, *M = ΔR+ − ΔR−*. Dotted horizontal and vertical lines mark zero recall response in each population. Each point represents one gene: yellow, genes with significant positive *M* that have stronger or more positive *ΔR*s in engram than non-engram neurons; purple, genes with significant negative *M*; gray, non-significant. Top three genes in each direction that are furthest from the diagonal are labeled, indicating maturation of state-dependent regulation. **d)** Modulation captures heterogeneous signals beyond recall response. Cross-classification of genes by their directions in *ΔR* in non-engram cells (y-axis) and engram cells (x-axis). Numbers indicate gene counts in each category. Many genes (n = 500 at s-value < 0.05, |LFC| > 0.1) show no significant *ΔR* overall but differential responses between engram and non-engram cells, demonstrating that *M* reveals signals missed by standard recall comparisons. **e)** Engram modulation is distinct between 7 d and 28 d. Scatter plot of *M_7_* (x-axis) and *M_28_* (y-axis) log_2_ fold changes. Each point represents one gene: yellow, genes with significant positive *M_28_*; purple, genes with significant negative *M_28_*; gray, non-significant for *M_28_*; black outlines, genes significant for both *M_28_* and *C*. **f)** Example genes illustrating engram modulation. *ΔR* estimates (log_2_ fold change, x-axis) for representative genes (y-axis) at 7 d and 28 d (facet) in engram (green) and non-engram (gray) cells. Error bars, ± standard error. *App* (Amyloid precursor protein) shows recall responsiveness that is selectively modulated in engram cells at 28 d, while *Adam10* (α-secretase) responds similarly in both populations. **g)** Functional enrichment across *ΔN*, *ΔR*, and *M*. Gene set enrichment of plasticity-related effectors and regulators across timepoint-aggregated *ΔN*, *ΔR*, *M* and timepoint-specific *M_7_* and *M_28_*, separated by direction of regulation (columns, Methods). Rows, functional gene sets. Colormap, fold enrichment. * FDR < 0.05; ** FDR < 0.01; *** FDR < 0.001. Effector programs are enriched in both *ΔR* and *M*, but regulatory processes are preferentially enriched in *M*, indicating selective recruitment of regulators in engram-modulated genes. **h)** Intrinsic plasticity genes are preferentially represented among down-modulated engram programs, suggesting that memory retrieval is accompanied by selective attenuation of excitability-related transcriptional responses. Fold enrichment (y-axis) of genes annotated as regulators of intrinsic neuronal plasticity was calculated separately among upregulated DEGs (top) and downregulated DEGs in recall response (*ΔR*), 7-day engram modulation (*M_7_*), and 28-day engram modulation (*M_28_*). Filled beige circles indicate significant enrichment, whereas open circles indicate non-significant enrichment. Circle size indicates -log_10_(FDR): * FDR < 0.05; ** FDR < 0.01; *** FDR < 0.001. **i)** Engram neurons selectively reshape recall-induced expression of ion-channel genes that control membrane excitability. Genes are shown on the y-axis, and the x-axis shows the recall-induced log_2_ fold change, *ΔR*, comparing recall with no-recall. Results are shown separately at 7 days (left) and 28 days (right) after learning. Green triangles indicate recall responses in engram neurons (*ΔR+*), gray circles those in non-engram (*ΔR−*). The vertical dashed lines mark no recall-induced change (log2 fold change = 0). Shown are the top three genes with the most positive and three with the most negative LFC values in *M*. *, FDR < 0.05; **, FDR < 0.01; ***, FDR < 0.001; ****, FDR < 0.0001. Colored asterisks on individual data points indicate significance for *ΔR*, and black asterisks on brackets indicate significance of *M* between a pair of *ΔR+* and *ΔR-*. **j)** Recall responses of voltage-gated channel genes are less dispersed in engram neurons, consistent with coordinated constraint of intrinsic excitability upon recall. The y-axis shows the recall-induced log_2_ fold change for each voltage-gated channel gene, and the x-axis shows recall responses in non-engram and engram neurons at 7 days (*ΔR−_7_* and *ΔR+_7_*) and 28 days (*ΔR−_28_* and *ΔR+_28_*) after learning. Each point represents one voltage-gated channel gene; gray points denote non-engram responses and green points denote engram responses. Lines connect the non-engram and engram estimates for the same gene within each time point, showing whether engram status facilitates or attenuates its recall response. The horizontal dashed line marks no recall-induced change. Selected genes with high response LFC magnitudes in *ΔR-_28_*, namely *Kcnab2* and *Kcnk2*, are labeled. Compared with non-engram neurons, voltage-gated channel genes exhibit significantly reduced variance in recall responses within engram neurons at both 7 days (*Var(ΔR−) = 0.089*, *Var(ΔR+) =* 0.054, variance ratio = 0.60, Pitman-Morgan test *P =* 0.008) and 28 days (*Var(ΔR−) = 0.110*, *Var(ΔR+) =* 0.075, variance ratio = 0.68, *P =* 0.046), indicating a compression of channel-gene response amplitudes in engram cells. Voltage-gated channel genes in Fig. 2i**,j** were defined by GO:0005244 (voltage-gated monoatomic ion channel activity) and further filtered for localization at neuronal cell body, dendrite, or initial segment of axon.

Specifically, we defined the ‘<u>n</u>o-recall engram’ (Δ*N*) signal as representing the differences between engram and non-engram neurons in the absence of memory recall. We defined the ‘recall response’ (Δ*R*) to denote the differences between recall and no recall conditions, capturing the effects of memory retrieval. The ‘engram <u>m</u>odulation’ (*M* = ΔΔ*R*) denotes the difference in Δ*R* between engram and non-engram neurons, capturing how the memory retrieval effects differ between the two cell sets. The ‘modulation <u>c</u>onsolidation’ (*C* = ΔΔΔ*R*) is the temporal shift in *M* between 7 and 28 days, capturing how the differences in the recall responses of engram and non-engram cells evolve over time (**Supplementary Fig. 3a**; **Methods**). In other words, *C* quantifies the extent to which the engram-dependent input-output relationship changes over the ∼4-week period when mPFC engram cells become necessary and sufficient for memory expression^3,4^. An advantage of using these 4 signals to analyze the complex, high-dimensional data from multiome sequencing is that it explicitly separates activity-, state-, and time-dependent regulatory effects.

The no-recall differences (Δ*N*) between engram and non-engram cells were modest, revealing 62 up- and 46 downregulated differentially expressed genes (DEGs) (|LFC|>0.1, s-value<0.05, **Fig. 2a**, **Supplementary Fig. 3b,c**). These DEGs included those linked to adhesion, gene regulation, and excitability, such as *Pcdh15*, *Nr6a1*, and *Ano3*, respectively. By comparison, when we aggregated recall responses across time points and engram states, Δ*R* revealed 957 up- and 903 downregulated DEGs; this included the upregulation of activity- and plasticity-associated genes such as *Tet3*, *Ntrk2*, and *Camkv*, as well as the downregulation of excitability-related genes such as *Scn1a* (**Fig. 2a**).

Engram modulation (*M*) revealed 555 up- and 626 downregulated DEGs, an order of magnitude more DEGs than identified by Δ*N*. Genes upregulated in *M* included *Cop1*, an E3 ubiquitin ligase targeting activity-regulated TFs including AP-1^41,42^, whereas the downregulated genes included *Rbfox1* and *Auts2*, which are implicated in RNA processing and neurite regulation. The upregulation of *Cop1* in *M* suggests engram cells may actively constrain recall-induced transcription by shortening the effective lifetime of activity-regulated transcription factors via the modulation of ubiquitination, thereby limiting recall-evoked downstream gene expression.

The modulation consolidation (*C* = *M*_28_ – *M*_7_) signal identified a smaller set of 22 up- and 42 downregulated DEGs (**Fig. 2a**). Strikingly, the top DEGs contributing to *C* exhibited a sign flip between *M*_7_ and *M*_28_ (**Supplementary Fig. 3c**). This reversal shows that consolidation does not simply strengthen or weaken the differential recall responses of engram *vs.* non-engram cells but can actually reverse whether a gene is preferentially engaged or attenuated in engram cells at recent *vs.* remote time points. In other words, this reversal indicates there is a wholesale exchange of roles over time in how engram and non-engram cells respond transcriptionally to memory recall.

Together, these results show that the transcriptional differences between engram and non-engram cells after a memory recall event were much larger than those without recall. This indicates that the engram cell state is characterized less by a constitutive transcriptional signature at rest and more by a broad transcriptional change that is only revealed at memory reactivation. Importantly, this conclusion was not dependent on the choice of a particular statistical threshold, since *M* remained larger than *ΔN* across a range of s-value cutoffs and was evident from the full distributions of effect sizes (**Supplementary Fig. 3d**). To this point, *M* showed more DEGs as well as log2-fold changes of greater magnitudes than *ΔN* (**Fig. 2a**, **Supplementary Fig. 3b**).

Although Δ*R* was substantially larger than Δ*N*, much of the recall-evoked transcriptional responses were common to engram and non-engram neurons (**Supplementary Fig. 3e**). We next examined how recall responses in engram and non-engram cells were related to one another. To do this, we compared Δ*R* in engram (Δ*R*+) and non-engram (Δ*R*-) neurons at each time point (**Fig. 2c**). In general, Δ*R* values of most genes were highly correlated between engram and non-engram cells.

However, a substantial subset of genes exhibited differences across the two cell sets, revealing an engram state-dependent modulation of transcriptional responses (*M*) at both 7 d and 28 d.

Notably, many of the genes revealed by *M* were undetected by a simpler analysis in which we averaged the Δ*R* results across time and engram state. Specifically, 500 DEGs identified by the *M* signal were not significant in the averaged Δ*R* signal (**Fig. 2d**). These included DEGs with induction or repression of recall-evoked responses selectively in engram cells, diminution in engram cells of shared recall-evoked responses, and polarity inversions in which memory recall regulated genes in opposite directions for engram *vs.* non-engram neurons (**Fig. 2d**, **Supplementary Fig. 3f**). Thus, *M* captured heterogeneous forms of engram-state-dependent transcriptional regulation that were obscured when recall responses were analyzed without accounting for engram state. This is biologically important, because it implies that memory recall did not elicit a uniform, activity-regulated transcriptional program across all neurons. Instead, the cells’ prior memory-encoding states reshaped the transcriptional responses to recall, allowing the same behavioral stimulus to engage different genes in engram and non-engram neurons of the same cell type.

We next asked how the structure of modulation itself changed over weeks. Although *M*_7_ and *M*_28_ showed comparable numbers of DEGs and effect sizes (**Fig. 2c, Supplementary Fig. 3f-h**), the genes identified by *M*_7_ and *M*_28_ were distinct (**Fig. 2e**). The top up-modulated genes in *M*_7_ were enriched for RNA splicing and local translation, whereas *M*_28_ included upregulation of transcriptional repressors (*Brinp1, Ifrd1*) and downregulation of structural plasticity genes (*Tmod1, Dctn1*; **Fig. 2c**). Overall, between 7 and 28 days, there was widespread remodeling of the engram-cell-specific modulation of recall-evoked responses.

The DEGs revealed by *C* (*i.e.*, *M*_28_ – *M*_7_) are those genes for which the passage of time led to a significant modulation of their engram-cell-dependent recall-evoked expression levels. These included regulators of RNA processing, such as *Srsf5*, and of synaptic signaling or vesicle release-associated genes, such as *Nrg2* and *Tspoap1*. Notably, this time-dependent modulation was significant for the *App* (Amyloid precursor protein) gene, of which recall-dependent upregulation was selectively attenuated in engram cells at 28 d but not 7 d. By contrast, *Adam10*, the gene for α-secretase, had recall responses that were similar across engram states (**Fig. 2f**). Although widely known for its involvement in Alzheimer’s disease, *App* is activity-regulated under physiological conditions and modulates synaptic plasticity^43–45^, suggesting that how plasticity-associated genes respond to recall in engram neurons is progressively reshaped over weeks.

### Engram cell modulation preferentially recruits regulatory programs

We next asked whether the DEGs we identified in the Δ*N*, Δ*R*, and *M* signals were associated with distinct functional programs. To do this, we compared the results for effector genes, which directly implement neuroplasticity, such as by altering synaptic function, neuronal structure, or membrane excitability, to those for regulator genes, which instead control other genes at the transcriptional or post-transcriptional levels. To classify genes as effectors or regulators, we used curated GO and Reactome term collections^46,47^ (**Methods**).

Plasticity-related effector genes were enriched in Δ*R* and *M*, whereas regulatory genes were preferentially enriched in *M*, including genes for chromatin regulation, RNA splicing, local translation, and protein ubiquitination (**Fig. 2g**). In accord, many up-modulated transcription factor (TF) genes in *M* were those implicated in chromatin state regulation, whereas many down-modulated TFs in *M* were those involved in activity-dependent transcriptional programs and cell identity (**Supplementary Fig. 3i**). Unexpectedly, the most distinctive signatures of engram modulation (*M*) were not expression changes for effector genes that implement neuroplasticity but instead were those of regulatory genes that control gene expression, RNA processing, protein translation and turnover. Thus, rather than reflecting only the execution of plasticity, *M* captured a higher-order molecular circuit that reshapes how engram cells deploy recall-evoked transcriptional programs.

Within the plasticity effector gene sets, Δ*N* had weaker enrichment for synaptic plasticity programs than Δ*R* or *M* and was instead enriched for genes affecting structural plasticity or neuronal membrane excitability. In other words, the no-recall engram (Δ*N*) differences emphasized plasticity that unfolds over slower time scales, including structural remodeling and excitability modulation, rather than acute forms of synaptic plasticity that arise just after memory encoding.

Notably, plasticity genes affecting excitability, including voltage-gated ion channels and other regulators of membrane excitability, showed a distinct regulatory pattern from that of other plasticity-associated effectors. While the genes affecting excitability were enriched among the upregulated *ΔR* genes, they were also strongly enriched among the downregulated *M* genes at both 7 and 28 days (**Fig. 2h**). This means that, in comparison to non-engram cells, by 4 weeks after memory encoding the engram cells had broadly attenuated recall-induced transcriptional programs that alter intrinsic excitability.

For example, recall responses were enhanced in engram neurons for *Hcn1*, *Grin1*, and *Kcnk2*, but attenuated for *Kcnab2*, *Kcnj6*, and *Cacna1d* (**Fig. 2i**). These genes include both excitatory and inhibitory regulators of membrane excitability, suggesting engram modulation does not simply bias ion channel expression toward a more or less excitable state. Instead, voltage-gated channel genes as a class showed a reduced range of recall responses in engram cells at both 7 d and 28 d (**Fig. 2j**), reflecting a compression of transcriptional response amplitudes across different ion channels.

Thus, while memory recall broadly activated genes affecting intrinsic excitability, the dynamic range of expression for these genes was diminished in engram cells. This shows there is diminished capacity for plasticity within consolidated engram states, potentially stabilizing previously modified circuits against excessive reconfiguration during recall and restricting memory interference^48,49^. Together, these findings suggest that cortical memory consolidation involves persistent regulation, not only of a static transcriptional state but also of transcriptional responses associated with memory recall. In other words, differential gene regulation itself is subject to state-dependent regulation.

### Engram neurons developed a distal chromatin engram signature over four weeks

Given that a cell’s engram state (engram or non-engram) modulated recall-evoked transcription (**Fig. 2a,c,f**), we asked whether the engram state is associated with distinct chromatin features, especially at distal candidate cis-regulatory elements (cCREs) including enhancers. To avoid biasing our results from chromatin accessibility changes near transcription start sites (TSSs) that might reflect consequences of transcription, we restricted our analysis of ATAC features to distal peaks (≥ 2 kb from any TSS). With this restriction, we computed LSI embeddings and found that the top reduced dimensions were dominated by cell-type effects (similar to **Fig. 1e**).

To quantify changes in chromatin accessibility space that were state-dependent and distributed across peaks, we adopted a decoding approach inspired by those commonly used in systems neuroscience to extract information from the activity traces of neural populations^50^. Here, our decoders were trained to identify signal directions in the high-dimensional ATAC LSI or gene expression PC space that distinguished between the 8 different nuclei classes in our experiment.

Starting with the glutamatergic cell types, we centered the embeddings of each neuron class to account for cell-type-specific effects, since for this analysis we sought to focus on state-, time-, and recall-dependent regulatory effects (**Methods**). We created multivariate logistic decoders that were trained to classify either chromatin accessibility or gene expression patterns across all glutamatergic cell types for the 8 different classes of nuclei, which we specified using linear combinations of the Δ*R*, Δ*N*, *M*, and *C* conditions (**Methods**). For the chromatin decoder, we used only accessibility peaks passing certain quality metrics (**Methods**). The resulting weights from both decoders are interpretable as regulatory axes and provide logit values that describe the regulatory effects of memory recall and engram state at either 7 or 28 days (**Fig. 3a, Supplementary Fig. 4a, Supplementary Table 2**).

**Figure 3.**
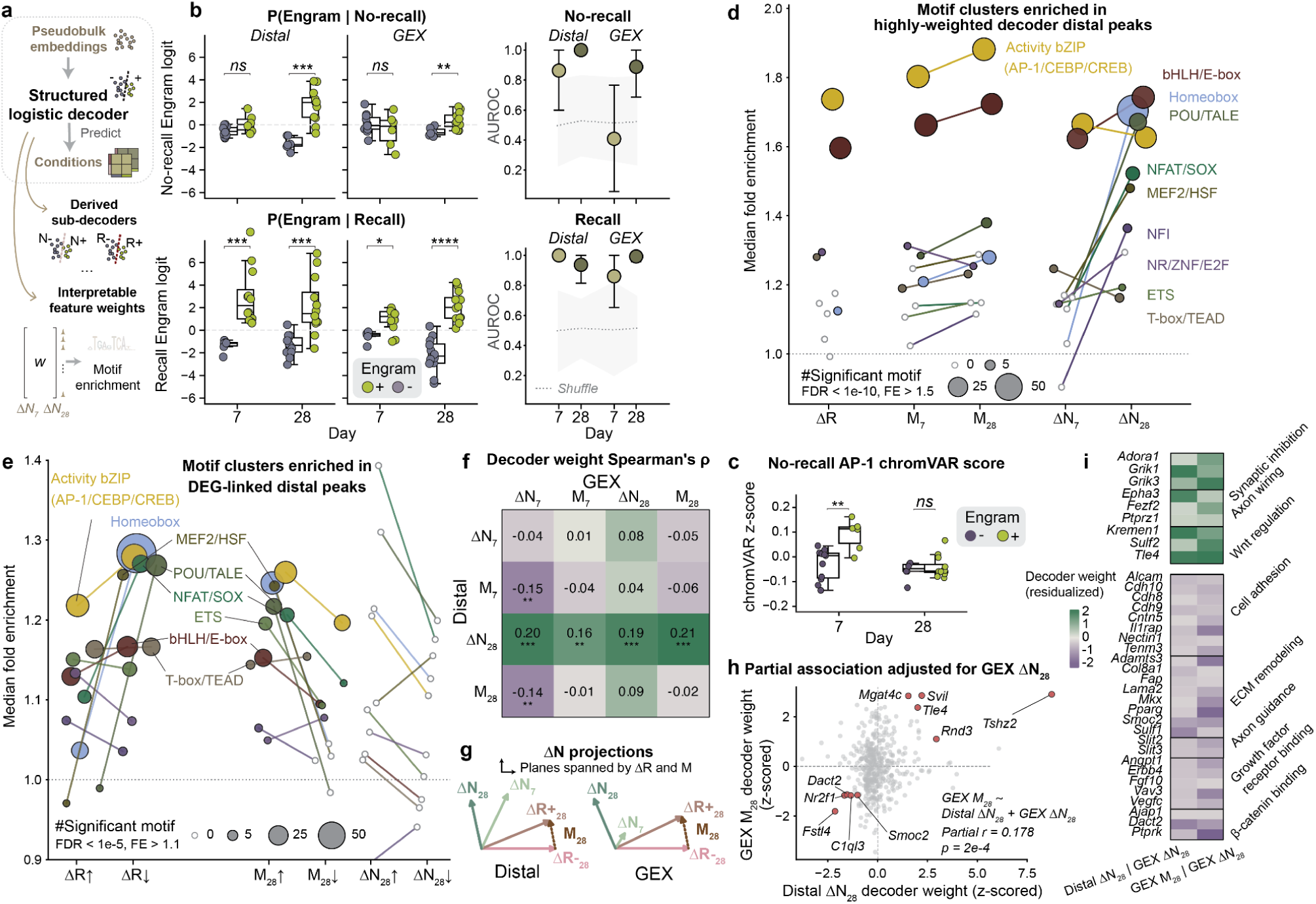
Decoders revealed a persistent distal chromatin engram signature that emerged over consolidation, stronger than no-recall transcription, and aligned with recall modulation. **a)** Decoders extract condition-specific regulatory directions from high-dimensional chromatin and transcriptome data. Distal ATAC features (≥2 kb from annotated transcription start sites (TSSs)) in glutamatergic neurons were embedded by latent semantic indexing (LSI) and residualized for cell-type effects. A constrained eight-class logistic regression decoder was trained with leave-one-experiment-out cross-validation to predict Time (7 d *vs.* 28 d), Engram (-/+), and Recall (N/R) states from distal ATAC embeddings (“Distal”) or gene expression (“GEX”) embeddings. **b)** At 28 d no-recall, engram and non-engram separate strongly in chromatin but weakly in RNA. Engram decoder logits (y-axes) conditioned on no-recall (top) and on recall (R+/R-, bottom) are shown for distal (ATAC) and GEX (RNA) modalities (columns). X-axis, 7 and 28 days. Distal chromatin separates engram from non-engram at 28 d no-recall but not 7 d no-recall, while GEX shows only weak separation at 28 d no-recall. At recall, both modalities separate robustly. Logit scores were calculated from held-out cross-validation folds (Methods). Right, decoder performance shown as AUROC (y-axis, mean ± 95% bootstrapped CI; gray band, shuffled control). X-axis, 7 and 28 days, Distal vs GEX modalities. Points represent pseudobulks (green, engram; gray, non-engram). Wilcoxon signed rank test: ns, p > 0.05; *, p ≤ 0.05; **, p ≤ 0.01; ***, p ≤ 0.001. **c)** The 28 d chromatin signature could not be explained by residual activity-dependent AP-1 activity. AP-1 chromVAR z-scores at no-recall showed higher AP-1 activity in engram cells compared to non-engram cells at 7 d but not 28 d, opposite to the maturation pattern of the distal engram signal. Dots represent pseudobulks: green, engram; gray, non-engram. X-axis, 7 d and 28 d. Y-axis, chromVAR z-score of AP-1 (JUN::FOS) motif. **d)** The 28 d chromatin engram signature is enriched for distinct regulatory TF motifs more commonly studied in development. Motif clusters enriched in distal peaks with robustly positive N+/N- sub-decoder weights. Motifs with FDR < 0.001 were grouped by PWM similarity; motifs with FDR < 1e-10 and fold enrichment (FE) > 1.5 were considered as significant (**Methods**). X-axis, decoder weight directions corresponding to *ΔN*, *ΔR*, or *M*, from which peak weights were extracted: *ΔR* aggregated across time points and engram states; *M_7_* and *M_28_*, *ΔN_7_*and *ΔN_28_*. Y-axis, median fold enrichment of each motif cluster, relative to a GC- and accessibility-matched background. Lines connect the same motif cluster between 7 d and 28 d within each decoder weight direction. Dot color, TF cluster identity; dot size, number of significant motifs per cluster; empty gray circles, clusters without significant motifs. **e)** The same TF motif grammar links chromatin state to recall-responsive and engram modulated genes. Motif enrichment of distal peaks linked by genomic proximity to *ΔR*, *M_28_*, and *ΔN_28_* DEGs. Motif clusters are defined as in Fig. 3d. X-axis, DEGs defined by regulatory quantity and direction: aggregated *ΔR*, *M_28_* and *ΔN_28_*, split into up- and downregulated DEGs. Y-axis, median fold enrichment of each motif cluster. Significant motifs were defined as FDR < 1e-5 and FE > 1.1. Dot color, TF cluster identity; dot size, number of significant motifs per cluster; empty gray circles, clusters without significant motifs. **f)** The 28 d no-recall chromatin signature aligns with RNA recall modulation. Spearman’s correlations between distal and GEX decoder weights show association between distal *ΔN_28_*and GEX *M_28_* (colormap and numbers). Distal peaks were linked to nearest expressed genes by genomic proximity, and peak weights were aggregated to gene-level scores. Analysis was restricted to 433 genes shared between distal and GEX decoders, with correlations computed in gene space. Rows, distal decoder weight directions; columns, GEX decoder weight directions. *, p ≤ 0.05; **, p ≤ 0.01; ***, p ≤ 0.001. **g)** 7 d no-recall chromatin signature is closer to recall response, 28 d signature closer to engram modulation. *ΔN_7_* aligns more with *ΔR+*, while *ΔN_28_* aligns more with *M* than with *ΔR+*, suggesting modulation inherits structure from the 28 d no-recall chromatin state. *ΔR+_28_*, *ΔR-_28_*, and *M_28_*, original geometry in decoder weight vector space; *ΔN_7_* and *ΔN_28_*, projections in the plane spanned by 28 d *ΔR+*, *ΔR-*, and *M*. Left, Distal decoder; right, GEX decoder. **h)** The chromatin-modulation link persisted after accounting for no-recall transcription. Residualized association between z-scored distal *ΔN_28_* and GEX *M_28_* decoder weights after regressing out GEX *ΔN_28_*. Top and bottom 5 genes ranked by concordance were highlighted in red. X-axis, distal *ΔN_28_* decoder weight aggregated by linked gene, z-scored residuals after regressing out GEX *ΔN_28_*; y-axis, GEX *M_28_* decoder weights, z-scored residuals after regressing out GEX *ΔN_28_*. **i)** Genes linking chromatin state to modulation were enriched for synaptic inhibition and Wnt signaling. Selected GO terms enriched among genes with high positive or negative concordance between residualized distal *ΔN_28_* and GEX *M_28_*. Rows, genes. Columns & hues: residualized decoder weights.

In the absence of memory recall, the gene expression decoder revealed no significant differences between engram and non-engram cells at 7 d and only weak differences at 28 d, consistent with the modest set of DEGs found earlier in *ΔN* (**Fig. 2a**). However, this was not the case for the chromatin accessibility decoder, which readily distinguished engram from non-engram cells at 28 d, thereby revealing a chromatin signature of engram neurons that emerged over the four weeks after fear conditioning (**Fig. 3b**). This notable difference, such that chromatin accessibility but not gene expression patterns readily identified the engram state at 28 d, indicates there is a long-lasting chromatin state that is hidden when looking solely at RNA expression. This general observation is broadly consistent with findings from bulk ATAC-seq and bulk RNA profiling studies of hippocampal engram neurons at 5 d after fear conditioning^14^.

In nuclei from mice that had undergone memory recall, both chromatin accessibility and gene expression decoders readily differentiated the engram cells at both 7 d and 28 d (**Fig. 3b**). The extent to which these findings differ from those above for the no-recall state highlights the strong effects of memory recall on both chromatin structure and gene expression and shows how recall unmasks chromatin changes that are nearly indiscernible in RNA expression patterns in the absence of recall.

Were the chromatin changes found for engram cells at 28 d in the no-recall condition reflective of the activity-dependent expression patterns and TFs induced at earlier time points, or did the chromatin state at 28 d reflect the engagement of a distinct set of TFs? To address this question, we first looked at the genome-wide level of chromatin accessibility for the AP-1 transcription factor family (comprising the FOS and JUN families) using the AP-1 chromVAR score^51^.

In engram cells, AP-1 chromVAR scores increased with memory recall, indicating that, as expected for cells we had trapped via *Fos* activation, memory recall elevates AP-1 activity (**Fig. 3b, Supplementary Fig. 4b**). However, without recall, at 7 d the engram cells showed higher AP-1 chromVAR scores than non-engram cells, but by 28 d after fear conditioning the engram and non-engram cells were no longer distinct in this regard. This indicates that the chromatin signature at 28 d in the no-recall state was not simply a reflection of AP-1 activity.

To identify which alternative programs were responsible for this chromatin signature in the absence of memory recall, we performed a motif enrichment analysis on the chromatin distal peaks that had positive decoder weights for Δ*N* at either 7 d (denoted Δ*N*_7_) or 28 d (Δ*N*_28_) (**Methods**). For example, the decoder peaks for Δ*N*_7_ were more accessible in engram than non-engram cells at 7 d.

Δ*N*_7_ peaks were enriched for activity-related bZIP TF family motifs (AP-1/CREB/CEBP), resembling those found in an analogous analysis for *ΔR* and *M* (**Fig. 3d**). Strikingly, however, for *ΔN_28_*a distinct motif grammar emerged, featuring chromatin motifs that are conventionally associated with development, cell lineage, and other durable programs for setting cell-states, instead of motifs associated with neural activity (**Fig. 3d**).

Motif clusters enriched for *ΔN_28_* included TFs from homeobox, POU/TALE, NFAT/SOX, MEF2/HSF, NFI, NR, ETS, and T-box/TEAD transcription factor families. We called these persistent TFs, abbreviated here as P-TFs. For example, the canonical OCT::SOX motif was enriched in *ΔN_28_*, and the joint OCT::SOX + AP-1 motif was enriched in both *M_28_* and *ΔN_28_* (**Supplementary Fig. 4c**). We also identified TFs with chromatin motifs that were significantly enriched in *ΔN_28_* decoder peaks and that had significant RNA expression changes captured in *ΔR* or *M* (**Supplementary Fig. 4d**). This concordance across chromatin and RNA analyses confirmed that, at least for a subset of the enriched chromatin motifs seen at 28 d, the enrichment was also mirrored in the RNA levels of the corresponding P-TFs.

In summary, the *ΔN_28_*chromatin signature, which distinguishes engram from non-engram cells at 28 d in the absence of memory recall, likely reflects a durable regulatory state of engram cells that involves developmentally associated P-TF programs instead of AP-1 related programs.

### Persistent transcription factors (P-TFs) are linked to engram-modulated genes

We next asked whether P-TFs are functionally linked to gene regulation. We mapped the DEGs identified above in the Δ*R*, *M*, and Δ*N* signals (**Fig. 2a**) to their corresponding distal chromatin peaks and then performed a motif enrichment analysis (**Methods**). The distal peaks linked to DEGs found in the *ΔR* and *M* signals were enriched in both activity-dependent bZIP (such as AP-1 and CREB) and P-TF motifs; this was especially the case for distal peaks for genes that were either downregulated in ΔR or upregulated in *M* (**Fig. 3e**). In line with the relatively small number of DEGs found in *ΔN* (**Fig. 2a**), few motifs were significantly enriched for *ΔN* (**Supplementary Fig. 4e**). Overall, separately from the chromatin data, the gene expression data pointed to developmentally related P-TFs as regulating memory recall responses as well as the engram-state-dependent modulation of these responses.

We next tested the idea that if P-TF-enriched *ΔN_28_* chromatin signatures prime the engram cells’ distinctive patterns of gene expression at memory recall, then the *M_28_* patterns of gene expression should correlate with *ΔN_28_* distal peak patterns. To test this hypothesis, we sought to determine whether the *M_28_* and *ΔN_28_*decoders shared a similar structure. We first mapped distal peak decoder weights to the space of RNA expression levels by assigning the values of the decoder peak weights to the expressed genes located nearest to each peak. By mapping the chromatin decoder to gene space in this way, we were able to compute a correlation coefficient between the two types of decoders, characterizing the extent to which they shared a common form (**Methods**). *M_28_* for gene expression was positively correlated with *ΔN_28_* for the distal peaks (Spearman’s rho = 0.21, p-value = 1.05e-5) but not with the distal-peak *M_28_* (rho = –0.02, p-value = 0.67; **Fig. 3f**; **Supplementary Fig. 4f**).

Further, for both distal peak and gene expression decoders, *ΔN_7_*was more similar to *ΔR+_28_*, but, as expected, *ΔN_28_*more closely resembled *M_28_* (**Fig. 3g**). In other words, at 7 days, the no-recall distal peaks and gene expression profiles resembled those reactivated by remote memory recall, whereas by 28 d these profiles were reshaped to reflect a different set of recall-evoked programs. We illustrated the geometry of this progression by projecting *ΔN* decoder weight vectors onto the plane spanned by *ΔR+*, *ΔR-*, and their difference *M* (**Fig. 3g**).

Genes for which chromatin accessibility and RNA levels were both substantially upregulated were revealed via the correlations between *ΔN_28_* for distal peaks and *M_28_* for gene expression (**Fig. 3h, Supplementary Fig. 4g**). This gene set was enriched for factors involved in synaptic inhibition, axon wiring, and Wnt signaling (**Fig. 3h,i**). The upregulation of *Adora1*, *Grik1*, and *Grik3*, from the top enriched GO term for ‘negative regulation of glutamatergic transmission’, suggests that by 4 weeks after memory encoding mature engram neurons have likely shifted toward a more synaptically restrained state, with reduced capacity for recall-evoked plasticity. On the other hand, the gene set with downregulation of both chromatin accessibility and RNA levels was enriched for genes involved in structural plasticity, including cell adhesion, ECM remodeling, and axon guidance factors, as well as growth factor receptor and β-catenin binding (**Fig. 3i**).

Collectively, these results suggest that by 28 days after memory encoding, a standing chromatin state in engram cells enables a down-modulation of recall-evoked programs associated with structural plasticity and an up-modulation of those for synaptic inhibition. In sum, these findings point to the existence of a chromatin-based mechanism for memory maintenance.

### Joint involvement of developmental and adult chromatin regulatory elements (cCREs)

We next asked whether the chromatin accessibility sites identified in *ΔN_28_*, which were enriched in engram cells for P-TF binding motifs, were reusing regulatory elements known to be active in brain development. To examine this, we applied our distal peak decoder, trained on the glutamatergic cell data from our fear conditioning study, to a mouse embryonic cerebral cortex scATAC-seq dataset^52^. Distal peaks at days E13.5, E15.5, and E18.5 in this published dataset overlapped with those identified by our distal peak decoder, and the percentage of distal peaks in the embryonic brain matching those in our adult brain dataset rose from ∼20% to ∼40% over E13.5 to E18.5 (**Supplementary Fig. 5a**; **Methods**). Across the three embryonic time points, there were 4,911 distal peaks that matched those used by our distal peak decoder. There were also 5,250 ‘adult-only’ distal peaks used by our decoder that were not found in the embryonic brain (**Fig. 4a**).

**Figure 4.**
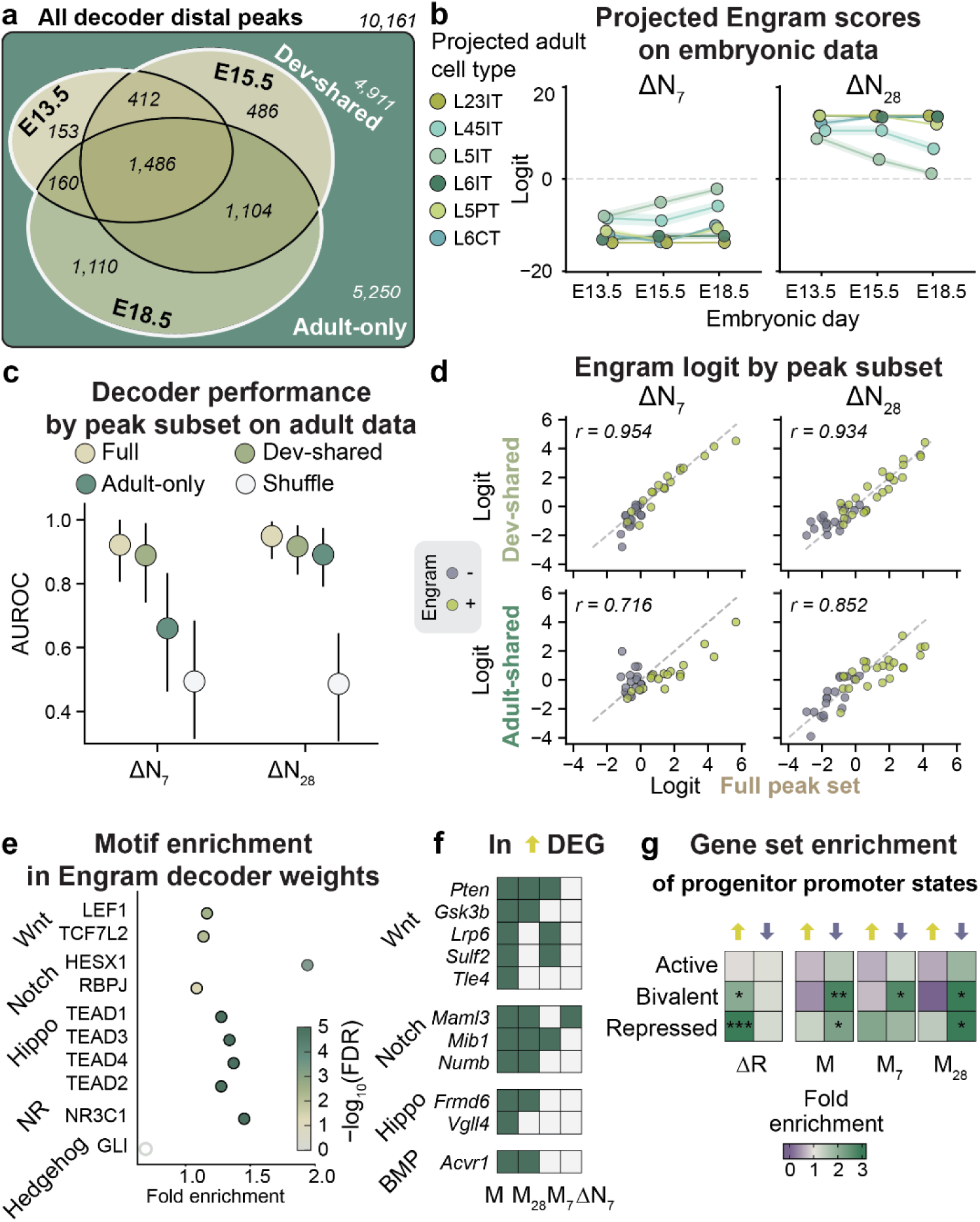
Persistent chromatin information is redundantly distributed across developmental and adult regulatory elements. **a)** Engram-associated distal peaks span both developmental and adult regulatory landscapes. Venn diagram of distal peaks used in the decoder, categorized by overlap with embryonic cortex datasets (E13.5, E15.5, E18.5). Peaks present in at least one embryonic time point were defined as “dev-shared” (n=4,911); remaining peaks were classified as “adult-only” (n=5,250). **b)** Embryonic cells resemble the mature (28 d) chromatin engram signature more than the early (7 d) one. Developmental scATAC-seq cell types were first annotated by label transfer from adult data, and glutamatergic cells from each embryonic day and cell type were aggregated into pseudobulks for decoder projection. Engram decoder scores projected onto developmental datasets show consistently negative *ΔN_7_*and positive *ΔN_28_* logits across embryonic days. Colors indicate glutamatergic cell types. X-axis, embryonic day; y-axis, projected no-recall engram decoder logit; shaded ribbons, standard errors over cross-validation folds. **c)** Developmental and adult regulatory elements independently support engram decoding. Decoder performance (AUROC) on adult data using the full peak set or subsets restricted to dev-shared or adult-only peaks. Performance is only modestly reduced when using subsets, indicating redundant encoding of engram-related information. X-axis, time points; y-axis, AUROC for *ΔN*. Tan, full peak set; light green, dev-shared peaks; dark green, adult-only peaks; white, shuffle control. **d)** Engram-related chromatin signals are preserved across developmental and adult peak subsets. Pseudobulk engram logits computed from dev-shared or adult-only peaks (x-axes) correlate strongly with logits derived from the full peak set (y-axis), with slightly stronger concordance for dev-shared peaks. Green, engram pseudobulks; gray, non-engram pseudobulks. **e)** The mature engram distal chromatin signature is enriched for motifs linked to developmental signaling pathways. TF motifs enriched in *ΔN_28_* engram decoder weights are associated with Wnt, Notch, Hippo, and nuclear receptor signaling pathways. X-axis, fold enrichment; y-axis, motifs grouped by pathway. Colors indicate -log_10_(FDR). **f)** Developmental signaling pathway genes are selectively engaged in no-recall and modulatory engram programs. Genes associated with Wnt, Notch, Hippo, and BMP signaling are upregulated in *M* and *ΔN* DEG sets. Rows, genes grouped by signaling pathway; columns, DEG sets including aggregated sets for *M, M_28_, M_7_*, and *ΔN_7_* using upregulated genes. Dark green, significant in the indicated DEG set; light gray, not significant. **g)** Engram-modulated genes are enriched for bivalent developmental promoter states. Gene set enrichment of neural progenitor promoter states across *ΔR* (aggregated over time points and engram states), *M* (aggregated over time points), *M_7_*, and *M_28_*DEG sets. Genes with bivalent or repressed promoter states are enriched in DEGs upregulated in *ΔR* or downregulated in *M*. Rows: active promoters, intermediate-CpG-density with H3K4me3; bivalent promoters, high-CpG-density with H3K4me3 and H3K27me3; repressed promoters, high-CpG-density with H3K27me3. Columns are separated by DEG direction. Color scale indicates fold enrichment. * FDR < 0.05; ** FDR < 0.01; *** FDR < 0.001.

We next asked whether neurons in the developing brain more closely resembled engram cells than non-engram neurons. By projecting our trained chromatin decoder onto the accessibility data for glutamatergic neuron types in this developmental dataset, we found that *ΔN_7_* logits were negative and those for *ΔN_28_* were positive at all time points, implying embryonic cells showed more of the chromatin signatures from the 28 d set of engram cells than those from 7 d (**Fig. 3a,b**, **Fig. 4b**; **Methods**). The *M_7_* and *M_28_* projected logits were more variable across embryonic cell types (**Supplementary Fig. 5b**). Notably, engram cells had not fully reverted to an embryonic-like state, as shown by the strong retention of chromatin accessibility peaks found only in the adult brain in *ΔN_28_* (**Supplementary Fig. 5c**).

For our adult brain data, decoders that were trained selectively on the subsets of accessibility peaks that were either shared across the adult and embryonic datasets or were present only in the adult data still classified engram cell states well, using similar decoder structures and with only a slight performance drop relative to decoders trained on the full set of accessibility peaks (**Fig. 4c,d**). These analyses suggest that developmental and adult cCREs carry engram-specific chromatin information redundantly.

Given the enrichment of the Wnt pathway (**Fig. 3i**), we further asked if developmental signaling pathways are involved along with developmentally-associated P-TF motifs and cCRE grammar. We found an enrichment of TF motifs related to Wnt, Notch, Hippo, and nuclear receptor pathways in the decoder weights (**Fig. 4e**), and an engram-specific modulation of genes in the Wnt, Notch, Hippo, and BMP pathways (**Fig. 4f**). The Hippo pathway enrichment was especially significant, suggesting its involvement in regulating cytoskeleton mechanisms^53^.

We next asked what chromatin regulatory states were associated with *ΔR* and *M* genes. Using neural progenitor promoter-state annotations, we found little enrichment for active H3K4me3-marked promoters but significant enrichment for bivalent and repressed promoter classes (**Fig. 4g**). Genes upregulated by recall (*ΔR* up) were enriched for repressed progenitor promoter states, consistent with genes that remain silent yet inducible under appropriate regulatory context^54,55^. Conversely, genes down-modulated in *M* were enriched for bivalent and repressed progenitor promoter states, with the enrichment for repressed states becoming significant at 28 d. Because *M* reflects the difference in recall response (*ΔR*) between engram and non-engram cells, this down-modulation suggests an attenuated induction of recall gene expression response. Thus, recall-induced genes that were attenuated in engram cells were preferentially associated with progenitor bivalent or repressive promoter states, suggesting that promoter-based chromatin mechanisms may contribute to selective gating of the engram recall response.

### GABAergic neurons enact distinct memory-related gene regulatory strategies

We next asked whether the transcriptional programs found in glutamatergic neurons were shared with GABAergic cells and found that the latter exhibited a markedly different regulatory architecture. Unlike glutamatergic neurons, in which the recall response (*ΔR*) and engram modulation (*M*) were prominent and differences between engram and non-engram cells in the absence of recall (*ΔN*) were modest, GABAergic cells exhibited more prominent (*ΔN*) features but weaker recall-evoked modulations (**Fig. 5a-b**). Thus, excitatory and inhibitory engram neurons use distinct strategies, distributing their memory-related transcriptional changes across different levels of regulation.

**Figure 5.**
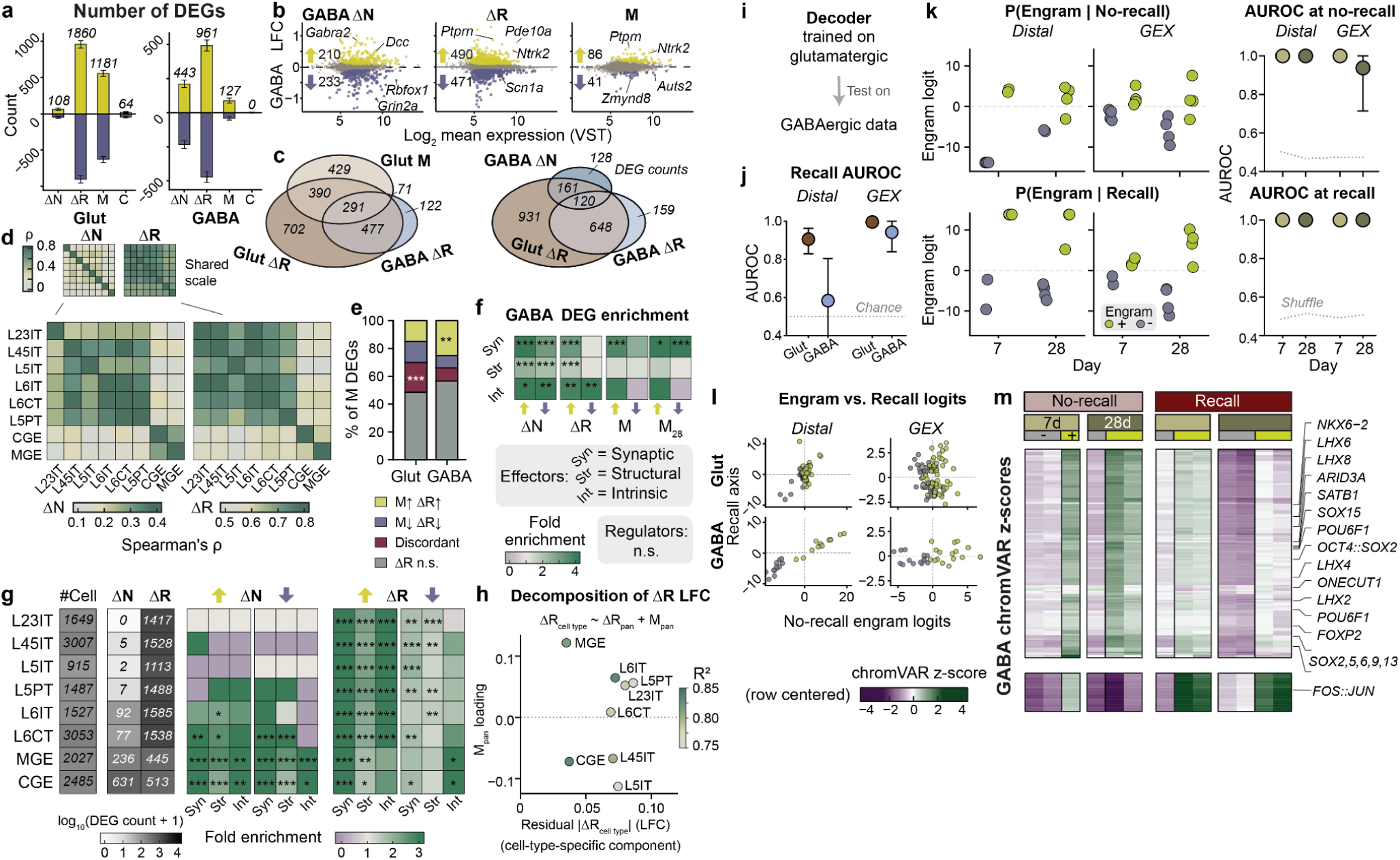
Glutamatergic and GABAergic neurons implement distinct regulatory strategies, with GABAergic programs being simpler, more evident at no-recall, and less modulatory. **a)** GABAergic neurons exhibit stronger no-recall programs and weaker engram modulation than glutamatergic neurons. Number of differentially expressed genes (DEGs; y-axis) for *ΔN* (no-recall engram), *ΔR* (recall response), *M* (engram modulation), and *C* (modulation consolidation) (x-axis) in glutamatergic (“Glut”) and GABAergic (“GABA”) neurons (facets), aggregated across time points and engram states. Yellow, upregulated; dark purple, downregulated. **b)** MA plots of *ΔN*, *ΔR*, and *M* in GABAergic neurons. Arrows indicate numbers of up- and down-regulated DEGs. Top three DEGs with the lowest s-values are labeled. X-axis, mean expression level (log_2_ VST); y-axis, log_2_ fold change (LFC) estimated by DESeq2. **c)** Recall-responsive transcriptional programs are largely shared between glutamatergic and GABAergic neurons. Venn diagrams showing overlap of DEG sets. Left, intersections among Glut *ΔR*, GABA *ΔR*, and Glut *M* highlight strong sharing of recall-responsive genes. Right, intersections among Glut *ΔR*, GABA *ΔR*, and GABA *ΔN*. **d)** No-recall engram programs are more cell-type-specific than recall response programs. Pairwise similarity of *ΔN* and *ΔR* programs across cortical neuronal cell types. Heatmaps show Spearman’s ρ between cell-type-specific *ΔN* or *ΔR* LFC vectors. Top, shared color scale; bottom, separate color scales for *ΔN* and *ΔR*. Rows and columns correspond to cell types. **e)** GABAergic modulation largely follows the direction of the recall response, with limited independent regulation. Concordance between *M* and *ΔR* DEGs. Categories: yellow, concordant up (upregulated in both *M* and *ΔR*); purple, concordant down (downregulated in both); maroon, discordant (opposite directions); gray, not significant in *ΔR*. Fisher’s exact test comparing Glut and GABA DEG counts: **, p < 0.01; ***, p < 0.001. Bars are normalized to 100%. **f)** GABAergic DEGs are enriched for plasticity effector programs but not regulator categories. Functional enrichment of GABAergic DEGs. Heatmap shows fold enrichment for synaptic (syn), structural (str), and intrinsic (int) plasticity-related gene sets as defined in Fig. 2. No significant enrichment was observed for regulator categories. Rows, functional gene sets; columns, directional DEG sets for each regulatory quantity. **g)** Cortical cell types span a gradient of regulatory strategies. Left, cell-type-resolved DEG counts for *ΔN* (aggregated over time points) and *ΔR* (aggregated over time points and engram states). Right, functional enrichment of plasticity effector gene sets across cell types. Heatmaps show log_10_(#DEG + 1) (left) and −log₁₀(FDR) for synaptic, structural, and intrinsic categories (right). **h)** Cell-type variations in *ΔR* are largely explained by shared recall and modulation programs rather than independent cell-type-specific programs. Decomposition of per-cell-type *ΔR* into shared *ΔR* and *M* components. Each point represents one cell type. X-axis, residual *|ΔR_cell type_|* (cell-type-specific component after removing pan-cell-class *ΔR* component); y-axis, loading on *M* (alignment with engram modulation). Color indicates model fit (R^2^). **i)** Decoder transfer analysis schematics. Distal ATAC and GEX RNA decoders trained on glutamatergic neurons (Fig. 3) were tested on GABAergic neurons. Performance was assessed on held-out Glut folds and the full GABA dataset. **j)** Recall-related signals transfer more robustly across cell classes in RNA than in chromatin. Recall decoding performance (AUROC; y-axis) for distal and GEX decoders (x-axis) in Glut and GABA neurons. Error bars, 95% bootstrapped confidence interval. **k)** Engram-related signals transfer robustly from glutamatergic to GABAergic neurons, despite weaker modulation in the latter. Engram classification in GABAergic neurons. Left, no-recall (N+/N−) and recall (R+/R−) logits (y-axis) across time points (x-axis); right, AUROC for distal and GEX decoders. Error bars, 95% bootstrapped confidence intervals. gray dotted line, shuffle control. **l)** Relationship between the no-recall engram axis and the recall axis. The recall axis (y-axis) was defined by Gram-Schmidt orthogonalization relative to the engram axis (**Methods**). X-axis, no-recall engram logits. Points represent biological pseudobulks (green, engram; gray, non-engram). **m)** GABAergic engram cells engage non-canonical TF motif programs similar to those observed in glutamatergic neurons (Fig. 3d**,f**). k-means clustering (k=5) of chromVAR z-scores of GABAergic neurons; only clusters with silhouette score > 0.45 are shown. Representative TF motifs are annotated. Rows, TF motifs; columns, experimental conditions.

Despite these differences in regulatory structure, recall-evoked transcription was broadly shared across excitatory and inhibitory cell classes. GABAergic *ΔR* genes showed extensive overlap with glutamatergic *ΔR* genes (**Fig. 5c**). Across all cell types, *ΔR* programs were more similar to one another than *ΔN* programs (**Fig. 5d, Supplementary Fig. 6a**). This indicates that while memory recall engages a shared program of neuronal activation, the engram state in the absence of recall was implemented using transcriptional configurations that were more cell-type-specific.

We next examined whether the genes identified by *M* in GABAergic cells represented a distinct modulatory program or mainly tracked the recall response. Compared with those found in glutamatergic cells, a larger fraction of GABAergic *M* genes were concordant with *ΔR*, and fewer showed discordant or *ΔR*-independent behavior (**Fig. 5e, Supplementary Fig. 6b,c**). This pattern suggests that GABAergic modulation appears to be more like a gain adjustment on recall-induced transcription. Functional enrichment analyses reinforced this interpretation. As in glutamatergic cells, GABAergic DEGs were enriched for plasticity effector gene sets across *ΔN*, *ΔR*, and *M* (**Fig. 5f**). However, unlike glutamatergic *M* genes, GABAergic DEG sets did not show significant enrichment for regulator categories (**Supplementary Fig. 6d**). Thus, transcriptional changes in inhibitory neurons provide less evidence for the regulator-heavy metaplastic programs found in excitatory neurons.

To determine whether this dichotomy between excitatory and inhibitory cells reflected a broad cell-class distinction or a continuum across neuronal subtypes, we quantified DEG counts and effector enrichment across cortical cell types. Cortical neurons exhibited a gradient of regulatory strategies, from modulation-heavy programs in upper-layer glutamatergic neurons to programs that were apparent in the absence of recall in deep-layer glutamatergic and GABAergic neurons (**Fig. 5g**).

We next asked whether these cell-type-specific recall responses represented independent transcriptional programs or could be explained by shared recall and modulation axes. Decomposing each cell type’s *ΔR* vector into shared recall and pan-modulation components revealed that cell-type variations in *ΔR* were largely captured by these common axes, with relatively few cell-type-specific components (**Fig. 5h, Supplementary Fig. 6e**). Overall, glutamatergic and GABAergic neurons use distinct regulatory strategies to differentiate engram from non-engram cells.

### Shared engram and recall axes generalize across neuronal cell classes

To test whether the low-dimensional chromatin and RNA axes identified in our analyses of glutamatergic neurons generalized to GABAergic neurons, we directly applied the decoders trained on glutamatergic neurons to the GABAergic data (**Fig. 5i**). Recall-related RNA signals transferred more robustly across cell classes than chromatin signals (**Fig. 5j**). Further, engram-related decoders generalized well, as both distal peak and gene expression decoders trained on glutamatergic cell data successfully differentiated the GABAergic engram and non-engram cell populations across both the no-recall and recall conditions (**Fig. 5k**). These results show that engram-related molecular axes are at least partly shared across excitatory and inhibitory neurons, even though specific genes may differ. Projection of GABAergic pseudobulk data onto no-recall engram and recall axes further showed that these axes were even more separable than those for glutamatergic neurons, in both distal chromatin and gene expression space (**Fig. 5l, Supplementary Fig. 6f**). ChromVAR analysis revealed GABAergic motif programs containing non-canonical developmental and cell-state-associated transcription factors (**Fig. 5m**). These motif programs resemble those observed in glutamatergic neurons. Altogether, these analyses revealed both shared and distinct molecular programs engaged by excitatory and inhibitory cortical neurons during memory consolidation.

## Discussion

Chromatin-based regulation may help remote memory circuits store information in a durable manner^3,14^. Here, we identify chromatin accessibility as a slow regulatory component that matures over weeks in cortical engram neurons and is predictive of how these cells will respond during memory recall events at future time points. The chromatin accessibility signature in engram cells was evident at distal candidate cis-regulatory elements and was also more prominent than engram signatures at the level of RNA profiles. Thus, our multiomic analyses suggest that chromatin accessibility not only provides a permissive backdrop for activity-dependent transcription but also appears to modulate an engram neuron’s basic input-output relationship dictating how future memory recall events will engage plasticity-associated gene programs.

The limitations of our study include its correlational design. Establishing causal roles for P-TFs, chromatin remodeling, and chromatin metaplasticity in memory processing will require perturbation experiments, such as P-TF knockdown, which we would predict to alter memory recall-dependent gene modulation (*M*). Further, activity-dependent genetic tagging is not a perfect means of labeling neurons involved in memory storage; some engram neurons may evade labeling, whereas other neurons that are not involved in storing the fear memory may be fluorescently labeled owing to unrelated electrical activation during the temporal window of genetic trapping. Thus, the neurons we trapped and analyzed should be regarded as a cell population that is highly enriched in but not identical to the full set of engram neurons within our mPFC tissue samples. As the exact relationship between *Fos* activation and a neuron’s electrical dynamics remains incompletely characterized in the live brain^5,28,56,57^, pooling all genetically trapped neurons together for data analyses may mask meaningful differences in electrical activation even among the population of trapped cells. Since our study focused on mPFC, future work will need to examine the extent to which our findings generalize to additional brain areas, forms of memory, and non-neural cell types.

### Chromatin metaplasticity as an efficient means of altering future neural responses

Neuronal plasticity constitutes an experience-dependent change in a neuron’s dynamical behavior or input/output function^58,59^. By comparison, metaplasticity is a higher-order form of plasticity, in which experience alters the cell’s ability to induce plasticity in the future, without necessarily changing the neuron’s present synaptic or electrophysiological properties^36,37^. This higher-order level of regulation reflects the multiple different ways in which a neuron can respond in parallel across multiple time scales to incoming electrical signals.

Namely, a single bout of electrical input can, in parallel, drive cellular and circuit level computations in the moment^50^, instruct changes in synaptic strength^59^, engage transcriptional programs, and also recruit metaplastic mechanisms that unfold over longer time periods^20,28^. Thus, gene expression evoked at memory recall is probably better viewed as part of the plasticity evoked by recall, such as activity-dependent strengthening or weakening of synapses^24,26,60,61^, rather than as part of the immediate expression of a memory at the seconds timescale. A learned change in such recall-evoked transcription is therefore metaplastic, quantified here as engram modulation (*M*). We refer to this chromatin-based regulation of future transcriptional responses as ‘chromatin metaplasticity’. This interpretation extends prior work showing that memory encoding increases enhancer accessibility, without corresponding transcriptional changes but with enhancer-promoter interactions reorganized at recall^14^. We similarly found that distal chromatin accessibility changes predict whether recall-evoked transcription is facilitated or attenuated, consistent with the idea that neuronal chromatin encodes metaplastic rules for future bouts of gene expression.

The metaplastic program that we identified was not merely an amplification or attenuation of plasticity. Notably, we found that glutamatergic neurons preferentially modulated regulator genes^62,63^, including those for chromatin regulation, RNA splicing, translational control, and protein turnover (**Fig. 2g**). The involvement of the ubiquitination system connects our results to past findings that long-term memory depends on protein degradation as well as protein synthesis^61,64–66^. More broadly, because proteostasis is energetically costly^67^, storing information as chromatin-encoded metaplastic rules that are engaged only during subsequent memory recall may be more energetically efficient than ongoing maintenance of memory-related gene expression.

Metaplasticity also helps resolve the inherent tension in the functional need to have circuits that are both stable and plastic, by selectively constraining plasticity rather than globally suppressing it^27,48,49,68^. This idea is related to the notion of memory allocation^69–71^, in which a neuron’s extant state, such as its levels of CREB-dependent excitability^69^ and chromatin plasticity^72^, influences whether the neuron is eligible to store a new memory. In the hippocampus, temporally proximal experiences are encoded by overlapping ensembles of neurons, whereas experiences that occur more distally in time are encoded by neural ensembles that are more distinct, thereby distinguishing the representations of unrelated events and reducing memory interference^73,74^.

These studies suggest that reducing interference between memories can be addressed, at least in part, by memory allocation mechanisms, allowing engram cells to remain available for memory recall but to avoid being overly repurposed for the storage of additional memories or plasticity events. Suggestive of a chromatin-based mechanism for such an allocation policy, engram cells preferentially up-modulated genes linked to downregulation of synaptic transmission (**Fig. 3i**) but attenuated recall-evoked expression of intrinsic excitability genes (**Fig. 2h,j**). Thus, chromatin metaplasticity may preserve established memories by limiting further plasticity in engram cells, while permitting neuronal activation for memory expression and new encoding elsewhere in the circuit.

### Cell-type-specific regulatory strategies

Prior single-cell omics studies have identified cell-type-specific molecular programs in memory, but these programs are usually described as gene lists or modules that are distinctive for particular cell classes^75,76^. By comparison, here we observed a more qualitative divergence between cell classes. Specifically, engram neurons of different cell types seemed to distribute their memory-related transcriptional changes across different levels of regulation.

In glutamatergic neurons, especially upper-layer IT neurons, engram and non-engram cells differed more in their modulation of recall responses (*M*) than in their no-recall expression levels (Δ*N*), whereas among GABAergic neurons, engram cells showed stronger no-recall transcriptional differences (Δ*N*) (**Fig. 2a**, **Fig. 5a,b,g**). This distinction may reflect the different circuit functions of the excitatory vs. inhibitory neuron classes. Excitatory neurons may preferentially encode memory-related changes as conditional input-output policies^77–79^, whereas inhibitory neurons may tune local circuit properties, such as levels of excitatory/inhibitory balance, via more sustained state changes^80,81^. Despite differences in specific genes and regulatory strategies, our decoders that were trained to distinguish glutamatergic engram cells transferred well to GABAergic neurons (**Fig. 5k**), suggesting that engram cell identity is represented along a shared low-dimensional regulatory axis^82,83^ that is robust to cell-type-specific molecular implementations (**Fig. 5l**).

### Reuse of developmental regulatory programs

P-TFs, as defined here, appear to be distinct from the activity-regulated TFs emphasized previously in the memory research literature^20,84^. Instead, P-TFs are associated with broad cell-state specification programs that are more commonly studied in the fields of developmental and cancer biology (**Fig. 3d,e**, **Fig. 4b**)^85^. Thus, our findings support the idea that aspects of gene regulation for development are reused in the adult brain in support of memory processing.

An evolutionary perspective suggests that such reuse may represent a neuronal specialization of a more ancient signal-responsive architecture that converts transient stimuli into persistent chromatin states. Unicellular holozoans already possess many components of this regulatory toolkit, including bZIP, bHLH, Fox, T-box^86^, SOX and POU TF families^87^. Among early-branching animals, sponges lack canonical neurons or muscles, yet coordinate tissue-level behaviors through Ca^2+^-dependent contractile and signaling systems. Downstream of these pathways, TFs can induce glutamate and GABA receptor subunits, cell fate regulators such as Srf and Fox, and adhesion genes^88,89^. Stimuli-activated MAPK signaling in sponges also primes chromatin accessibility at regions enriched for AP-1 and FOX motifs^90^. Vertebrate neurons deploy this same regulatory toolkit during late-phase plasticity through *FoxO6*, *Sox11* and AP-1^91^. Beyond the nervous system, AP-1-centered regulatory logic similarly establishes persistent chromatin states in non-genetic cancer drug resistance memory^92^ and inflammation memory^93^. In this view, Ca^2+^-dependent chromatin remodeling in memory may represent a neural adaptation of an ancient cellular plasticity program rather than a uniquely neural invention.

This perspective suggests that conserved developmental regulatory programs, reused in memory, may also provide a useful lens for interpreting disease-associated chromatin states. For instance, recent evidence suggests that schizophrenia-associated increases in chromatin accessibility in cortical neurons recapitulate fetal progenitor-like accessibility states^94^. Likewise, the functional effects of many neuropsychiatric disease risk variants remain hidden in resting states but are unmasked by neuronal activation^95^. Together, these observations suggest that disease might arise from the inappropriate deployment of evolutionarily conserved cellular regulatory programs connecting neuronal activation, gene expression, and chromatin states.

### Outlook

Our findings also motivate a broader conception of memory. For the past ∼70 years, memory has been viewed primarily in Hebbian terms as involving experience-dependent changes in synaptic strength^96–99^. Although synaptic plasticity is surely involved in many forms of memory, the extent to which synaptic weights alone are sufficient to specify a memory remains unclear^98,100–102^.

Our findings show not only that engram cells have a long-lasting, distinctive chromatin state from that of non-engram cells but also that this state shapes an engram cell’s transcriptional response to a future memory recall event. In other words, neuronal metaplasticity includes long-lasting changes to chromatin structure. This finding is mirrored by the inclusion of regulatory factors among the set of genes for which RNA expression at memory recall is distinct in engram cells. Thus, long after the initial storage of a memory, engram cells run their own distinct transcriptional programs at memory recall. Based on this key finding, we propose that the state of a neuron should be defined to include all variables that shape the cell’s response to future inputs. This includes not just synaptic weights and connectivity patterns but also the chromatin state, which determines the neuron’s responsiveness to its future inputs. The memory trace therefore includes the metaplastic effects arising from changes in chromatin architecture.

Testing this broader conception of memory will require experiments that go beyond taking snapshots of neuronal gene expression or chromatin accessibility patterns at different time points. Instead, future experiments will need to probe engram neurons’ altered transcriptional programs by providing neurophysiological or behaviorally driven inputs at time points that are well past the initial encoding of memory. By combining behavioral assays of memory with functional imaging of neural activity, activity-dependent genetic trapping, spatial multi-omics, and targeted perturbations of enhancers, promoters, or transcription factors, neuroscientists can now examine how chromatin state alterations affect neural circuit dynamics and memory performance^39,103–105^.

### Author contributions

Y.K., W.J.G. and M.J.S. conceived the study. Y.K. designed the experiments with input from W.J.G. and M.J.S. Y.K. and X.S. performed fear conditioning experiments. Y.K. performed the molecular experiments, generated the multiome datasets, developed and performed the computational analyses, and generated and edited the figures. Y.K. interpreted the results with input from W.J.G. and M.J.S. Y.K. wrote the manuscript. W.J.G. and M.J.S. edited the manuscript. All authors approved the final manuscript. W.J.G. and M.J.S. supervised the study.

## Acknowledgements

We thank Jordi Fernandez-Albert for early experimental work and advice on nuclei isolation and sorting, Jane Li for animal husbandry, Soon il Higashino and Yanping Zhang for experimental support, and Samuel H. Kim, Guiping Wang, Benjamin Parks, Immanuel Abdi, Betty B. Liu, Selin Jessa, Lacramioara Bintu, Michael Z. Lin, Will Allen, and Thomas Kenying Lau for helpful discussions. We thank the Stanford Shared FACS Facility (RRID:SCR_017788) for help with fluorescence-activated nuclei sorting and the Stanford Functional Genomics Facility (RRID:SCR_002050) for access to Chromium instrumentation and support of pilot experiments. This work was supported by the Carol & Gene Ludwig Family Foundation. M.J.S. acknowledges support from the Howard Hughes Medical Institute, the Stanford Knight Initiative for Brain Resilience, and a Vannevar Bush Faculty Fellowship from the U.S. Department of Defense. W.J.G. acknowledges support from the Arc Institute and the Chan Zuckerberg Biohub, and funding from the NIH (R01HG013317, DP1HG013599, R01NS128028).

## Methods

### Mice

All animal experiments were performed in accordance with protocols approved by the Stanford University Administrative Panel on Laboratory Animal Care. TRAP2 mice carrying the Fos2A-iCreERT2 allele (Jackson Laboratory, stock no. 030323) were crossed with CAG-Sun1/sfGFP reporter mice (Jackson Laboratory, stock no. 021039). Mice were maintained on a C57BL/6 background and group-housed, with a maximum of five mice per cage, under a 12-h light-dark cycle (lights on from 07:00 to 19:00), with food and water available ad libitum. Male and female mice aged 12–20 weeks were used. Mice were singly housed beginning 3 days before contextual fear conditioning and remained singly housed throughout the experiment. Mice were assigned to one of four experimental groups defined by time after learning (7 or 28 days) and recall exposure (no recall or recall).

### Reagents

#### 4-Hydroxytamoxifen

4-Hydroxytamoxifen (4-OHT; Sigma-Aldrich, cat. no. H6278) was first dissolved in ethanol with shaking at 60 °C for 10 min and then diluted with Kolliphor EL (Sigma-Aldrich, cat. no. C5135) to prepare 20 mg ml^−1^ stock solutions. Stock solutions were stored at −20 °C for up to several weeks. On the day of use, stock 4-OHT was diluted with an equal volume of PBS to a final concentration of 10 mg ml^−1^.

#### Anti-NeuN antibody

Anti-NeuN antibody (clone A60, Sigma-Aldrich, cat. no. MAB377) was conjugated to XFD647 NHS Ester (AAT Bioquest; chemically equivalent to Alexa Fluor 647 NHS Ester). 2 μl of 10 mg ml^−1^ dye in DMSO was added to 112.5 μl of 1 mg ml^−1^ antibody in 0.1 M sodium bicarbonate buffer and incubated at room temperature for 15 min. The conjugated antibody was purified by buffer exchange using Amicon Ultra 0.5 ml centrifugal filter units and resuspended in PBS containing 0.1% sodium azide at a final antibody concentration of 1 mg ml^−1^.

### Experimental design

For each biological replicate, mPFC tissue from a pooled group of mice was processed together, and nuclei were separated by fluorescence-activated sorting into GFP− and GFP+ populations. GFP− and GFP+ nuclei derived from the same mouse pool were processed as separate 10x Multiome libraries. Most biological replicates consisted of pooled tissue from four mice, with one replicate consisting of three mice, and were sex-balanced when possible. In total, 17 libraries spanning eight experimental conditions were generated; sample-level metadata are provided in **Supplementary Table 1**.

#### Contextual fear conditioning and genetic trapping

Mice were handled daily in a ventilated hood for 3 days before contextual fear conditioning. On the conditioning day, mice were staged in the hood for 15 min before transport to the behavioral room in opaque transfer boxes. All mice, including those assigned to recall and no-recall groups, received 10 mg ml^−1^ 4-hydroxytamoxifen (4-OHT; Sigma-Aldrich, cat. no. H6278) at 50 mg kg^−1^ via intraperitoneal injection immediately before conditioning. Contextual fear conditioning was performed in a chamber with a stainless-steel bar floor (Coulbourn Instruments), treated between animals with 70% ethanol and 1% Virkon-S disinfectant solution. Contextual cues, including lighting, background noise, transport procedures and handler, were kept constant across conditioning and recall sessions. Mice were allowed to explore the chamber for 120 s before receiving five 2-s foot shocks (0.75 mA) delivered at 30-s intervals. Mice remained in the chamber for 60 s after the final shock before being returned to their home cages. Each mouse underwent a single conditioning session. For recall experiments, mice were returned to the same context for 5 min either 7 or 28 days after conditioning. Recall sessions were initiated at 15-min intervals such that tissue from each mouse could be collected exactly 60 min after recall onset. No-recall control mice remained in their home cages and were collected at matched early-morning times.

Freezing behavior was recorded and quantified using FreezeFrame software (version 4; Coulbourn Instruments). Freezing was defined as an episode of immobility lasting longer than 1 s and was quantified across the entire conditioning or recall session using a fixed detection threshold applied uniformly across animals.

#### Nuclei isolation and fluorescence-activated nuclei sorting

Mice were euthanized by cervical dislocation, and brains were rapidly dissected and rinsed in ice-cold DPBS. To isolate medial prefrontal cortex tissue, the olfactory bulbs were removed and a 5-mm-thick coronal section was obtained using a mouse brain matrix. Bilateral diagonal cuts were made to remove ventrolateral tissue and enrich for medial prefrontal cortex. Tissue from multiple mice was pooled and homogenized on ice in a dounce homogenizer (Kimble Kontes; #885300-0002) containing 1 ml nuclei extraction buffer [250 mM sucrose, 25 mM KCl, 5 mM MgCl_2_, 20 mM HEPES-KOH, 65 mM β-glycerophosphate, 0.5% IGEPAL CA-630 (Sigma-Aldrich, cat. no. I8896), 1× cOmplete protease inhibitor (Sigma-Aldrich, cat. no. 11836170001), 1 mM DTT, 0.2 mM spermine, 0.5 mM spermidine, 60 U ml^−1^ RNasin Plus RNase inhibitor (Promega, cat. no. N2611), and 2.5% normal goat serum (Invitrogen, cat. no. 10000C)]. Tissue was homogenized with 20 strokes each of pestles A and B, incubated for 5 min at 4 °C with rotation, and filtered through a 40-μm Flowmi cell strainer (Sigma-Aldrich, cat. no. BAH136800040).

Filtered nuclei were stained with Alexa Fluor 647-conjugated anti-NeuN antibody (1:1,000) for 15 min at 4 °C and DAPI (20 μM) for 2 min. Nuclei were sorted on a BD FACSAria II directly into 5× nuclei buffer (10x Genomics nuclei buffer supplemented with 5 mM DTT and 5 U μl^−1^ Protector RNase Inhibitor; Sigma-Aldrich, cat. no. 3335399001). DAPI-positive nuclei were gated sequentially for singlets, NeuN^+^, and GFP expression (**Supplementary Fig. 1a**). Across FANS experiments with available gate statistics, NeuN+ nuclei represented 70.0 ± 3.0% of singlet nuclei (mean ± s.e.m.; n = 5 sorting dates), and GFP+ nuclei represented 0.73 ± 0.11% of NeuN+ singlet nuclei (n = 6 sorting dates). Sorted nuclei were processed immediately using the Chromium Single Cell Multiome ATAC + Gene Expression workflow.

#### Single-nucleus multiome library preparation and sequencing

Single-nucleus ATAC and gene expression libraries were prepared using the Chromium Single Cell Multiome ATAC + Gene Expression platform (10x Genomics, PN-1000283) according to the manufacturer’s protocol. Following fluorescence-activated nuclei sorting, nuclei were loaded onto Chromium Next GEM chips for droplet generation, barcoding and library preparation. Libraries were sequenced on an Illumina NovaSeq 6000 platform (Novogene) to the sequencing depth recommended by the manufacturer.

### Initial single-cell processing and quality control

#### Multiome alignment and initial cell calling

Sequencing reads were aligned to the mm10 mouse reference genome, and initial cell-associated barcodes were identified using the Cell Ranger ARC pipeline hosted on the 10x Genomics Cloud Analysis platform. The cloud-managed software version was not provided. Each library was processed independently using default parameters. No additional barcode rescue or manual modification of the initial cell calls was performed.

#### Modality-specific quality control

Single-nucleus ATAC-seq data were processed using ArchR (v1.0.2). Nuclei were retained based on the joint distribution of transcription start site enrichment and unique fragment counts. Thresholds were selected by visual inspection of sample-specific quality-control distributions to separate the high-quality nucleus population from low-quality barcodes. Nuclei were required to have a transcription start site enrichment score ≥5 and blacklist ratio <0.05, with additional fragment-count thresholds applied as needed on a per-sample basis. Chromosomes X, Y, and mitochondrial DNA were excluded from downstream ATAC analyses.

Single-nucleus RNA-seq data were processed using Seurat (v5.2.1). Nuclei were first required to have more than 200 RNA unique molecular identifier (UMI) counts before doublet detection. After doublet detection, nuclei were retained when they contained at least 500 and no more than 7,500 detected genes, at least 1,000 and no more than 80,000 RNA UMI counts, and less than 5% mitochondrial transcripts. RNA counts were log-normalized. 2,000 highly variable genes were selected using the mean–variance plot method, and expression values were centered and scaled.

Only barcodes passing both RNA and ATAC quality-control criteria and detected in both modalities were retained for subsequent joint analysis. A total of 35,396 nuclei passed ATAC quality control, 33,753 nuclei passed RNA quality control, and 28,389 nuclei were shared between the RNA and ATAC datasets.

#### Doublet detection and cell-type filtering

Doublet scores were calculated from the RNA count matrix using scDblFinder (v1.20.0) with cluster-aware doublet detection (clusters = TRUE, dbr.sd = 0) and an expected doublet rate of 0.006 per 1,000 cells (dbr.per1k = 0.006) separately within each sample. Joint filtering then excluded nuclei that were either non-neuronal, with low label-transfer confidence, or with TSS enrichment scores less than 5.25. Initial per-cell doublet classification removed 898 nuclei, leaving 27,491 singlets. Following initial clustering and annotation, residual doublet populations were identified using cluster-level scDblFinder scores. At clustering resolution 3, three clusters with median scDblFinder scores greater than 0.5 were removed, excluding 401 nuclei. After re-clustering at resolution 1, two additional small clusters with median scDblFinder scores greater than 0.5 were removed, excluding 139 nuclei. In total, approximately 1,438 nuclei were removed through per-cell and cluster-level doublet filtering.

#### Data exclusion

Two samples, YK01 and YK10, were excluded from differential expression analyses after review of sample-level quality-control metrics and integrated projection behavior. Both samples showed evidence of atypical sample structure, including low usable nuclei representation after filtering, reduced concordance with biological replicates in variance-stabilized pseudobulk expression profiles, and aberrant projection patterns in the ATAC UMAP, most prominently in L6CT neurons. Because differential expression analysis tests each gene individually, such sample-level deviations can have outsized effects on dispersion estimation, coefficient fitting, and gene-wise significance calls. We therefore excluded YK01 and YK10 from gene-level differential expression analyses.

All samples were retained for decoder analyses. Unlike differential expression analysis, the decoder analysis asks whether condition information is distributed across multivariate molecular profiles. Decoder inputs were regularized or dimension-reduced, and performance and operator contrasts were computed from held-out predictions across cross-validation folds. Thus, decoder analysis does not rely on gene-wise inference from individual features and is less sensitive to the influence of any single outlier sample. Including all samples in the decoder analysis provided a more conservative test of whether condition structure generalized across the full experimental dataset.

#### RNA and ATAC dimensionality reduction

RNA and ATAC dimensionality reduction and visualization were performed separately. For the RNA modality, principal-component analysis was performed using the 2,000 highly variable genes after expression scaling. The first 40 principal components were used to construct the nearest-neighbor graph and generate the RNA UMAP embedding. For the ATAC modality, iterative latent semantic indexing (LSI) was performed in ArchR using addIterativeLSI on the genome-wide TileMatrix with LSIMethod=2, using ArchR defaults of two iterations, 25,000 variable features, and 30 LSI dimensions. An ATAC UMAP embedding was generated from the first 30 LSI dimensions. UMAP visualizations were used for exploratory analysis and visualization only and were not used for statistical testing.

#### Cell type annotation

Cell type annotation was guided by an annotated Allen Institute mouse cortex single-cell RNA-seq reference dataset^106^. The reference dataset was restricted to prefrontal cortical regions, including MOs_FRP, ACA, PL-ILA-ORB, and AI, and non-neuronal cell classes were excluded. To reduce computational cost while preserving representation across cell types and anatomical regions, reference cells were downsampled by stratified sampling across region and subclass labels. Transfer anchors between the Allen reference and experimental RNA-seq data were identified using Seurat’s *FindTransferAnchors* function based on the top 2,000 highly variable genes and principal components 2–40. Cell *cluster*, *subclass*, *neighborhood*, and *class* annotations were transferred to individual nuclei using *TransferData*.

Cell clustering and annotation were performed in parallel. Transferred annotations were compared with Seurat clusters by cross-tabulation and combined with cluster-level marker gene expression and manual inspection of RNA and ATAC embeddings to assign final cell identities. Cluster annotations were subsequently propagated to the matched ATAC profiles. Populations corresponding to non-prefrontal cell types from neighboring anatomical regions were excluded from downstream analyses. Following annotation and filtering, the final dataset contained 17,669 high-quality mPFC neuronal nuclei.

#### Peak calling

Accessible chromatin peaks were identified using the ArchR reproducible peak-calling workflow, excluding chromosomes X and Y and mitochondrial DNA. Pseudobulk accessibility profiles were generated separately for each annotated cell type using addGroupCoverages, with a minimum of 20 cells per pseudobulk and up to 300 cells sampled per replicate. Multiple pseudobulk replicates were generated for each cell type (minimum 13, maximum 16 replicates). Peaks were called independently for each cell type using MACS2 through ArchR’s addReproduciblePeakSet function (groupBy = “cell_type”, reproducibility = “1”). Overlapping peak calls were merged using ArchR’s iterative reproducible peak-merging procedure to generate a unified peak set across all cell types. Peaks located on chromosome X or Y, or mitochondrial DNA were removed prior to construction of the final peak accessibility matrix. The resulting merged peak set contained 508,952 peaks and was used to generate the single-cell peak accessibility matrix with ArchR’s addPeakMatrix function.

#### Pseudobulk generation

Pseudobulk profiles were generated separately for the RNA and ATAC modalities by aggregating counts within each sample-by-cell-type combination. For RNA, raw UMI counts were summed across nuclei to generate pseudobulk gene-expression profiles. For ATAC, counts from the ArchR peak accessibility matrix were summed across nuclei to generate pseudobulk peak-accessibility profiles. Genes were retained for downstream RNA analyses if they had at least 50 counts in at least 25% of pseudobulks. ATAC peaks were retained if they had at least 10 counts in at least 5% of pseudobulk samples and were detected in at least two samples. To identify variable ATAC features for downstream dimensionality reduction and decoder analyses, size factors and dispersions were estimated using DESeq2. Peaks with evidence of excess dispersion relative to the fitted mean-dispersion trend (upper-tail χ² p-value < 0.1) were retained, yielding 15,382 variable peaks. Library-size normalization factors were estimated using the DESeq2 poscounts method, and variance-stabilizing transformation was applied to generate the feature matrix used for dimensionality reduction and classifier training.

#### Peak-to-gene assignment

To associate distal regulatory elements with candidate target genes, peaks were assigned to the nearest transcription start site (TSS) of expressed protein-coding genes. The expressed-gene universe was defined from the RNA pseudobulk dataset as genes with at least 20 counts in at least 25% of pseudobulk samples and detected in at least five samples.

Gene coordinates were obtained from the mm10 reference annotation. Peak-to-gene assignments were generated using GenomicRanges (v1.58.0) by calculating the distance from each peak center to the nearest TSS. Peak-gene pairs separated by more than 500 kb were excluded from downstream analyses.

#### ChromVAR analysis

Transcription factor motif activity was quantified using chromVAR deviation scores as implemented in ArchR, calculated from the single-cell ATAC-seq data and subsequently averaged within each sample-by-cell-type pseudobulk for downstream statistical analyses.

### Statistical framework and core analyses

#### Definition of regulatory quantities

Across differential expression and decoder analyses, we used a common NRMC framework to define regulatory quantities from condition-specific molecular profiles: no-recall engram state (*ΔN*) captures differences between engram and non-engram neurons in the absence of recall; recall response Δ*R* = Δ_R_*x*; engram modulation *M* = Δ*E*Δ*R* = Δ*E*Δ*_R_x*; modulation consolidation *C* = Δ*TM*= Δ*T*Δ*E*Δ*Rx*, where *R* ≡ *Recall*, *E* ≡ *Engram status* (engram vs. non-engram), *T* ≡ *Timepoint*, and *x* denotes gene expression or peak accessibility features.

#### Differential expression analysis

Differential expression analysis was performed using DESeq2 (v1.46.0) on pseudobulk RNA profiles. Glutamatergic and GABAergic neurons were analyzed separately to reduce model complexity and account for major transcriptional differences between neuronal classes. Within each neuronal class, a single generalized linear model was fit across all constituent cell types using the design formula: ∼ CellType + Time + Recall + Engram + Time:Engram + Time:Recall + Recall:Engram + CellType:Engram + CellType:Recall + Time:Recall:Engram where Time indicates the interval between learning and tissue collection (7 or 28 days), Recall indicates whether animals underwent memory recall before sacrifice, and Engram indicates TRAP-labeled (GFP+) versus unlabeled (GFP−) neurons. Cell type was included as a categorical covariate, and cell type-specific recall and engram effects were modeled through interaction terms. Reference levels were set to 28 days, no recall, GFP−, and L6CT (glutamatergic) or CGE (GABAergic).

Pseudobulks containing fewer than 50 nuclei were excluded prior to analysis. Two additional samples, YK01 and YK10, were excluded based on quality-control metrics (**Supplementary Fig. 1, Supplementary Methods**). Effect sizes were shrunk using adaptive shrinkage (ashr), and statistical significance was assessed using s-values, which estimate the posterior probability of an incorrect sign for the reported effect. Differentially expressed gene (DEG) sets were defined by s-value < 0.05 and |LFC| > 0.1.

#### Contrast definition for differential expression

The regulatory quantities introduced in **Fig. 2** were estimated as linear contrasts of coefficients from the fitted DESeq2 generalized linear model.

No-recall engram:

- *^ÄN^_28_* ^= Engram^_pan-cell-class_
- *ÄN_7_* = Engram_pan-cell-class_ + Time:Engram

Where 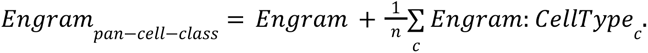

Recall response:

*ΔR-_28_* = ^Recall^_pan-cell-class_

*ΔR+_28_* = ^Recall^_pan-cell-class_ + Recall:Engram

*ΔR-_7_* = ^Recall^_pan-cell-class_ + Time:Recall

*ΔR+_7_* = ^Recall^_pan-cell-class_ + Recall:Engram + Time:Recall + Time:Recall:Engram

- *ÄR_pan_* = Recall_pan-cell-class_ + 0.5 * Recall:Engram + 0.5 * Time:Recall + 0.25 * Time:Recall:Engram

where 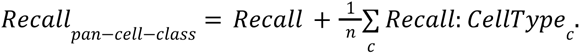

Engram modulation:

- *M_7_* = Recall:Engram + Time:Recall:Engram
- *M_28_* = Recall:Engram

Modulation consolidation:

- *C* = M_28_ − M_7_ = - Time:Recall:Engram

#### Functional gene set enrichment analysis

Gene set over-representation analysis was performed on differentially expressed genes corresponding to each regulatory quantity and direction using predefined gene sets and the set of expressed genes of a given broad cell class as the background universe. Enrichment was assessed using Fisher’s exact test, and p-values were adjusted for multiple testing using the Benjamini-Hochberg procedure. FDR < 0.05 was considered significant.

Gene Ontology (GO) annotations from the Biological Process, Molecular Function, and Cellular Component ontologies were used to construct curated gene-set collections representing broad classes of effector and regulatory functions. Related GO terms were grouped into higher-order categories using a YAML-based annotation framework, and enrichment was evaluated at the category level rather than for individual GO terms. Over-representation analysis was performed using Fisher’s exact test. GO annotations were obtained and processed using the R package clusterProfiler (v4.14.4).

Promoter chromatin-state gene sets were obtained from the Molecular Signatures Database (MSigDB)^107^. Three promoter-state annotations derived from genome-wide histone modification profiling in mouse neural progenitor cells were examined: MIKKELSEN_NPC_ICP_WITH_H3K4ME3 (active/permissive promoters), MIKKELSEN_NPC_HCP_WITH_H3K4ME3_AND_H3K27ME3 (bivalent/poised promoters), and MIKKELSEN_NPC_HCP_WITH_H3K27ME3 (repressed promoters). These gene sets were curated in MSigDB from the chromatin-state maps reported by Mikkelsen et al. and were used without modification.

### Predictive and comparative decoder analyses

#### Decoder model construction

To compare chromatin and transcriptional structure across experimental conditions, we trained constrained multinomial logistic regression decoders on pseudobulk profiles. Separate decoders were trained for ATAC and RNA modalities. For ATAC, features consisted of latent semantic indexing (LSI) components computed from distal peak (≥2 kb away from any TSS) accessibility profiles. For RNA, features consisted of principal components computed from pseudobulk gene-expression profiles. Before model fitting, features were first standardized, then centered by each cell type.

Rather than fitting each experimental condition independently, the decoder was parameterized using biologically interpretable component weight vectors. Condition-specific decoder weights were modeled as linear combinations of five component weight vectors: a pan-time recall effect (*ΔR*), the mean and temporal components of the no-recall engram effect (*ΔN_mean_* and *ΔN_time_*), and the mean and temporal components of engram-specific recall modulation (*M_mean_* and *M_time_*). Here, *ΔN_mean_* and *M_mean_* represent the average engram and modulation effects across 7 and 28 days, respectively, whereas *ΔN_time_* and *M_time_* represent their temporal differences between 7 and 28 days. Equivalently, *ΔN_7_ = ΔN_mean_ − ΔN_time_*, *ΔN_28_ = ΔN_mean_ + ΔN_time_*, *M_7_ = M_mean_ − M_time_*, and *M_28_ = M_mean_ + M_time_*. The fixed coefficient matrix used to reconstruct the eight condition weights is in **Supplementary Table 2**.

For each pseudobulk, logits were normalized only across the four classes from the corresponding time point. Thus, 7-day samples were evaluated against the four 7-day classes, and 28-day samples were evaluated against the four 28-day classes. Per-sample weights were set so that each of the eight experimental conditions contributed equally to the loss. The model was fit by minimizing the weighted masked cross-entropy loss with L2 regularization on the five component weight vectors. Intercepts were included for the non-reference classes, with 7N− and 28N− intercepts anchored to zero. The regularization strength was selected by cross-validated grid search.

Cross-validation was performed using leave-one-experiment-out fold splitting, where an experiment denotes one pool of animals processed together. In most cases, GFP+ and GFP− samples from the same animal pool were treated as the same experiment and assigned to the same fold. For this glutamatergic decoder, three experiments were the sole source of one experimental condition. Holding out each of these experiments in full would remove that condition from the training set, so these folds were split by glutamatergic cell type using a fixed partition: L2/3 IT, L4/5 IT, and L5 IT in one sub-fold, and L5 PT, L6 CT, and L6 IT in the other sub-fold. All other experiments were held out as intact folds. In total, 11 folds were used. Held-out predictions from cross-validation were used for downstream decoder analyses. Engram logits were computed within each time point and recall condition as the log-odds of GFP+ versus GFP− classes. Decoder performance was evaluated using AUROC for engram classification within no-recall and recall conditions. Null distributions were estimated by label shuffling, and uncertainty was estimated by bootstrap resampling.

Decoder-derived regulatory weight directions were obtained from the learned component weight vectors. For ATAC decoders, component weights learned in LSI space were projected back to distal peak space to estimate peak-level contributions to each regulatory direction. Peaks with consistently positive weights across cross-validation folds were identified using a one-sided test across folds followed by Benjamini-Hochberg correction. Stable positively weighted peaks were used for downstream motif enrichment analysis.

#### Motif enrichment analysis

Motif enrichment analysis was performed using ArchR motif annotations generated from the JASPAR 2022 motif database. For decoder-derived ATAC contrasts, foreground peak sets consisted of stable positively weighted distal peaks identified from the corresponding decoder component weights as described above. For differential expression-linked peak analyses, foreground sets were defined as the union of distal peaks assigned to genes significant for each regulatory quantity and direction.

For each foreground set, a matched background set was sampled from all called peaks after excluding the foreground peaks. Background peaks were matched to each foreground peak by GC content and accessibility, using 10 equal-width GC-content bins and 10 quantile-based accessibility-score bins. For each foreground peak, up to 2,000 candidate background peaks were sampled from the same GC/accessibility bin, and the union of sampled peaks across the foreground set was used as the matched background. GC content was calculated from the mm10 genome using BSgenome.Mmusculus.UCSC.mm10, and accessibility matching used the peak score stored in the ArchR peak set.

Motif enrichment was assessed using a hypergeometric test comparing motif occurrence in the foreground set against the matched background set. P-values were adjusted using the Benjamini-Hochberg method, and enriched motifs were defined as FDR < 1 × 10^−10^ and fold enrichment ≥ 1.5 for decoder-derived peak sets, or FDR < 1 × 10^−5^ and fold enrichment > 1.1 for DEG-linked peak sets.

#### Embryonic chromatin projection analysis

To compare embryonic chromatin accessibility with adult memory-related accessibility states, embryonic ATAC-seq peaks from E13.5, E15.5, and E18.5 were matched to the adult distal peak universe used for decoder analysis. Peaks were represented as 501 bp windows, and embryonic peaks overlapping adult peaks were considered matched. Single embryonic nuclei were projected into the adult distal-peak LSI space using the adult peak universe, inverse-document-frequency weights, and SVD components learned from adult pseudobulk ATAC profiles. Embryonic cells were assigned adult cell-type labels by k-nearest-neighbor label transfer in the adult LSI space. Adult single-cell ATAC profiles from this study were used as the reference. Both adult and embryonic cells were TF-IDF transformed using the adult IDF weights and projected onto the adult SVD components. For each embryonic cell, the 15 nearest adult cells were identified using cosine distance, and the projected cell type was assigned by majority vote. Downstream decoder projection was restricted to cells assigned to glutamatergic classes. Developmental pseudobulks were generated by summing embryonic peak counts within each embryonic time point and projected adult glutamatergic cell type. Pseudobulk profiles were TF-normalized, transformed using the adult IDF weights, projected onto the adult distal-peak LSI basis, and centered by the corresponding adult glutamatergic cell-type centering vector before decoder analysis.

Developmental pseudobulk profiles were then passed through the adult-trained 8-class multinomial decoder. To interpret embryonic profiles relative to adult memory states, decoder probabilities were normalized separately within the 7-day and 28-day adult reference frames using time-masked softmax normalization. Decoder-derived scores, including no-recall engram effect, recall response, and engram modulation of recall response, were computed as described above. These scores correspond to contrasts over recall, engram status, and time in the adult design framework.

#### Contribution of embryonically accessible and adult-specific peaks

To assess the contribution of developmentally shared and adult-specific distal regulatory elements to decoder performance, adult pseudobulk accessibility profiles were reprojected through the trained decoder after masking selected peak subsets. Masked features were set to zero prior to LSI projection and classification. Three feature sets were evaluated: the full distal peak set, a developmentally shared peak set, and an adult-specific peak set. For each feature set, pseudobulk samples were projected into the corresponding LSI representation and evaluated using the same cross-validation framework used for decoder training. Decoder performance was quantified using area under the receiver operating characteristic curve (AUROC) for Recall, Engram, *ΔN*, *ΔR*, and *M* contrasts. Confidence intervals were estimated by bootstrap resampling, and shuffled-label controls were used to establish null performance distributions.

---

**Supplementary Figure 1.**
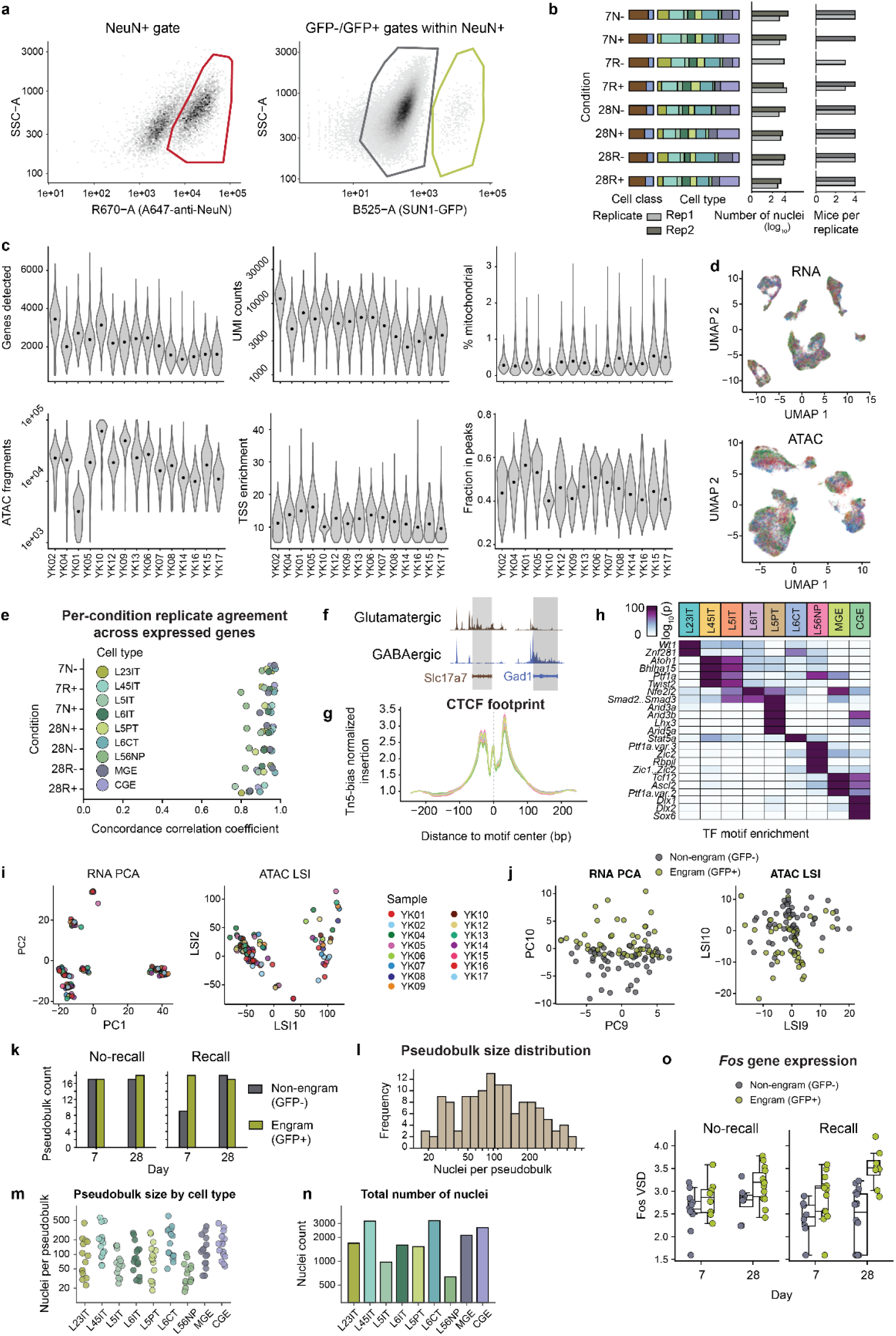
High-quality sorting and sequencing establish a robust mPFC single-nucleus multiome dataset. **a)** Representative FANS gating strategy. DAPI+ singlet nuclei were selected after debris and doublet exclusion, followed by isolation of NeuN+ neuronal nuclei and separation into SUN1-GFP− and SUN1-GFP+ gates. Left, side-scatter area (SSC-A; y-axis) versus Alexa Fluor 647-conjugated anti-NeuN fluorescence (R670-A; x-axis), with the NeuN+ gate outlined in red. Right, SSC-A (y-axis) versus SUN1-GFP fluorescence (B525-A; x-axis) within NeuN+ nuclei, with GFP− and GFP+ gates outlined in gray and green, respectively. **b)** Mouse, sample, and cell-composition accounting across the eight experimental conditions. Conditions are defined by time after learning (7 or 28 days), recall status (N, no-recall; R, recall), and GFP gate (− or +). Rows indicate condition. The stacked bars show the relative abundance of annotated cell classes and cell types within each condition; colors denote cell identity. Adjacent horizontal bars show the number of nuclei on a log_10_ scale and the number of mice contributing to each biological replicate. Replicates 1 and 2 are distinguished by dark- and light-gray shading, respectively. **c)** Per-nucleus RNA and ATAC quality-control metrics for each sample. Samples are shown on the x-axes. From left to right, top row: number of genes detected per nucleus, RNA unique molecular identifier counts, and percentage of RNA counts assigned to mitochondrial genes; bottom row: number of high-quality ATAC fragments, transcription start-site enrichment, and fraction of ATAC fragments overlapping called peaks. Violin widths represent the density of nuclei at each value, and black points indicate sample medians. **d)** RNA and ATAC UMAPs colored by sample. Each point denotes one single nucleus. Color legend is shown in panel **i)**. **e)** Agreement of gene-expression profiles between biological replicates. The y-axis indicates experimental condition, and the x-axis shows the concordance correlation coefficient between replicate pseudobulk RNA profiles for the same condition and cell type. Each point represents one cell type, with colors indicating the annotated cell type. **f)** Representative aggregate chromatin-accessibility tracks at canonical neuronal marker loci. Tracks show aggregate ATAC insertion coverage for glutamatergic and GABAergic neurons across genomic position at the *Slc17a7* and *Gad1* loci, respectively; shaded regions highlight cell-class-specific accessibility at marker gene bodies. **g)** Aggregate CTCF footprint across cell types. The x-axis indicates distance from the center of CTCF motif occurrences, and the y-axis shows Tn5-bias-normalized insertion frequency. Colored lines represent individual cell types. Depletion of insertions at the motif center, flanked by increased insertion frequency, indicates occupancy of CTCF-bound sites. **h)** Cell-type-specific enrichment of transcription-factor motifs in accessible chromatin. Columns indicate cell types, grouped into glutamatergic, GABAergic, and non-neuronal classes, and rows indicate transcription-factor motifs. Color represents motif-enrichment significance as −log_10_(*P*), with darker shading indicating stronger enrichment. **i)** First two reduced dimensions for RNA and ATAC encoding cell type differences. Each point denotes one pseudobulk and is colored by sample. **j)** The 9th and 10th reduced dimensions for RNA and ATAC encode condition-related cell states. Each point denotes one pseudobulk and is colored by engram (green) or non-engram (gray). **k)** Summary of pseudobulks retained for downstream analyses after requiring more than 15 nuclei per sample-by-cell-type pseudobulk. The x-axes indicate day after learning and the y-axes indicate the number of retained pseudobulks for no-recall and recall conditions, respectively; gray and green bars denote GFP− and GFP+ populations. **l)** Distribution of pseudobulk sizes, with nuclei per pseudobulk on the x-axis and pseudobulk frequency on the y-axis. **m)** Pseudobulk sizes per cell type. X-axis, cell type; y-axis, number of nuclei per pseudobulk. Colors indicate cell types as in **g)**. **n)** Total number of nuclei per cell type. X-axis, cell type; y-axis, total number of nuclei. Colors indicate cell types as in **g)**. **o)** *Fos* is expressed at higher levels in engram than non-engram cells especially at 28 d recall. Y-axes, DESeq2-normalized Fos expression level (VSD). Each point denotes one pseudobulk.

**Supplementary Figure 2.**
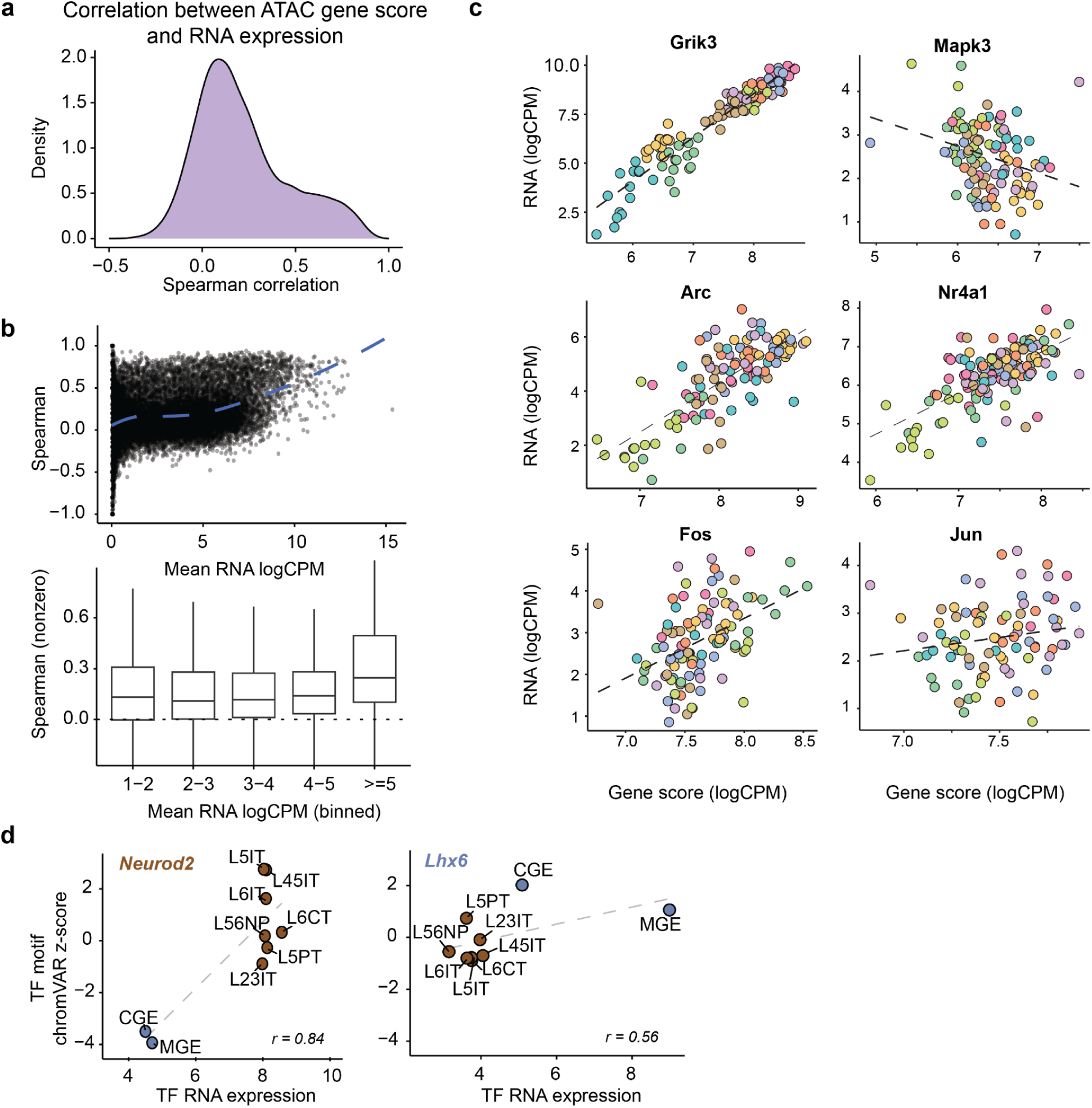
Cross-modality coupling between RNA expression and chromatin accessibility is highly gene dependent. **a)** Distribution of Spearman correlations between matched ATAC gene scores and RNA expression across genes. Most genes show modest correlation, while a subset of genes show positive cross-modality coupling. **b)** Gene-level RNA–ATAC coupling only modestly increases with expression levels. Top, Spearman correlation between gene score and RNA expression plotted against mean RNA abundance. Bottom, genes binned by mean RNA expression show only slightly stronger modality concordance among more highly expressed genes. **c)** Representative genes illustrating diverse RNA–ATAC relationships across nuclei. Canonical cell type markers (e.g. *Grik3*) show tight coupling, and housekeeping genes (e.g. *Mapk3*) show no or negative coupling. Immediate early genes (e.g. *Arc*, *Nr4a1*, *Fos*, and *Jun*) span a spectrum of high to low cross-modality coupling. *Grik3* and *Mapk3* were selected as the highest coupled and lowest coupled genes; *Arc*, *Nr4a1*, *Fos*, and *Jun* were selected as representative immediate early genes. Each dot represents one pseudobulk and is colored by cell type. **d)** TF activity inferred from chromVAR deviation scores is correlated with TF RNA expression across cell types for representative lineage-associated transcription factors, including *Neurod2* and *Lhx6*. Each point represents a cell type; labels indicate major neuronal subclasses.

**Supplementary Figure 3.**
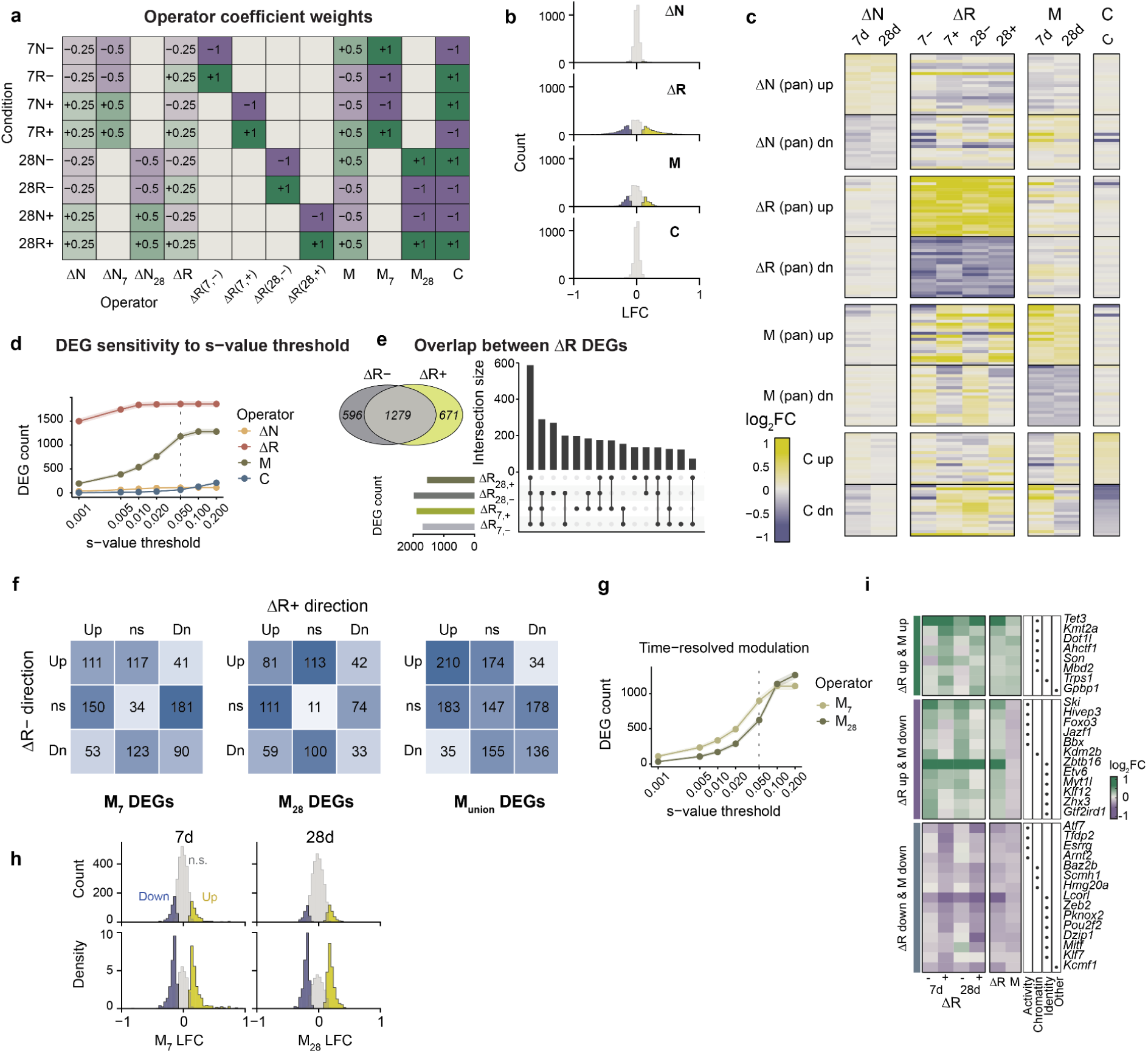
Operator-based differential expression reveals bidirectional engram modulation of recall responses. **a)** Linear contrast weights used to estimate coefficients from the eight experimental conditions (rows), including no-recall engram differences at 7 d and 28 d after learning (*ΔN_7_, ΔN_28_*), recall responses in each time point and gating condition (*ΔR*), time-resolved engram modulation of recall responses (M_7_, *M_28_*), their aggregation over time points (*M*), and the maturation of modulation across consolidation time (*C*). **b)** Distributions of log2 fold-change estimates (x-axis) for *ΔN, ΔR, M*, and *C*, with significant positive and negative effects highlighted. Y-axis, gene counts. Shaded ribbons indicate 95% Wilson confidence intervals for DEG counts. Yellow, upregulated; purple, downregulated; gray, non-significant. **c)** Heatmaps of log2 fold changes for genes grouped by significant up- or down-regulation for each regulatory quantity, showing how no-recall engram state, recall response, engram modulation, and maturation effects relate across time points. **d)** Sensitivity of DEG counts (y-axis) to s-value threshold (x-axis) for each regulatory quantity (color). Shaded ribbons indicate 95% Wilson confidence intervals for DEG counts. **e)** Overlap among recall-response DEGs across time point and gating (engram) conditions, with total *ΔR+* (recall response in engram cells) and *ΔR−* (recall response in non-engram cells) DEG counts summarized by Venn diagram. **f)** Relationship between underlying recall-response directions in engram and non-engram cells for engram modulation DEGs. Heatmaps show genes significant for *M_7_, M_28_*, or their aggregation over time points (*M*). Rows indicate the direction of the recall response in non-engram cells (*ΔR−*), and columns indicate the direction of the recall response in engram cells (*ΔR+*). “Up” and “Dn” indicate significant recall-induced up- or down-regulation, and “ns” indicates no significant recall response. Numbers indicate the number of modulation DEGs in each *ΔR−/ΔR+* category. Diagonal entries represent concordant recall responses between engram and non-engram cells, whereas off-diagonal entries represent gain, loss, attenuation, potentiation, or reversal of recall responses in engram cells. This classification reveals whether engram modulation reflects enhanced induction, reduced induction, selective induction, enhanced repression, reduced repression, or polarity switching. **g)** Sensitivity of time-resolved modulation DEG counts (y-axis) to s-value threshold (x-axis) for M_7_ and M_28_. **h)** Distribution of engram-modulation log2 fold changes (LFC) at 7 d and 28 d after learning. Histograms show the full distribution of *M_7_* and *M_28_* LFC across genes. Top row shows gene counts, and bottom row shows the same distributions normalized to density to better compare the shape of significant and non-significant effects. Yellow bars indicate genes with significant positive engram modulation, where recall-induced expression is higher in engram cells than in non-engram cells. Purple bars indicate genes with significant negative engram modulation. Gray bars indicate genes without significant modulation. Vertical lines mark the log2 fold-change positions of significant up- and down-modulated genes. Both time points show bidirectional modulation, with positive and negative interaction effects rather than a single global shift in recall responsiveness. **i)** TF gene expression patterns associated with representative regulatory categories. Heatmaps show log2 fold changes for transcription factor genes grouped by their differential expression patterns across recall response and engram modulation quantities. Rows are TF genes, and columns show recall-response coefficients at 7 d and 28 d in non-engram and engram cells (*ΔR−* and *ΔR+*), followed by engram modulation (*M*). Green indicates increased TF expression and purple indicates decreased TF expression. TFs are grouped into three illustrative patterns: genes with coordinated recall induction and positive modulation (*ΔR* up & *M* up), genes with recall induction but negative modulation (*ΔR* up & *M* down), and genes with recall repression and negative modulation (*ΔR* down & *M* down). The annotation bar on the right summarizes broad functional classes for each TF, including activity-associated, chromatin-related, identity-related, and other regulatory categories. Filled dots indicate membership in each functional annotation class.

**Supplementary Figure 4.**
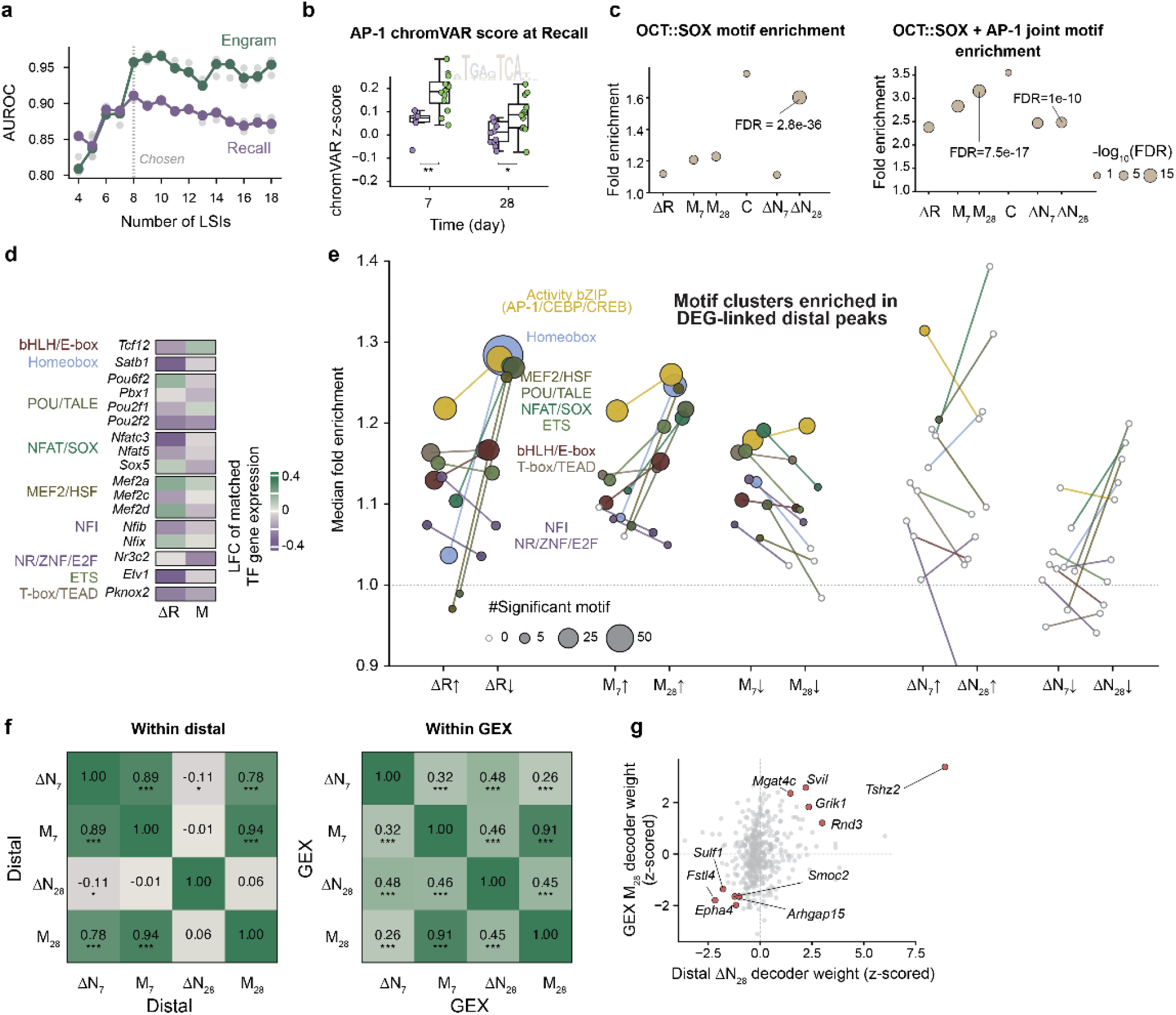
Decoder analysis links persistent chromatin engram signatures to motif programs and transcriptional modulation. **a)** AUROC as a function of the number of distal LSI dimensions included in the decoder grid search. Performance is shown as the median across five cross-validation folds for the engram decoder (green) and recall decoder (purple); non-median fold values are shown as gray dots. The selected number of LSI dimensions is indicated by the gray vertical dashed line. **b)** AP-1 motif activity at recall, measured by chromVAR z-score, stratified by time point and distal engram decoder score. Boxplots show AP-1 chromVAR scores in nuclei grouped by decoder-predicted chromatin engram state, showing higher AP-1 accessibility in engram cells compared to non-engram cells at 7 d and 28 d. **c)** The canonical OCT::SOX motif is enriched in no-recall engram at 28 d, and the OCT::SOX–AP-1 joint motif is enriched in both recall-time modulation and no-recall engram. Motif enrichment for OCT::SOX and OCT::SOX–AP-1 joint motifs across distal peak sets linked to *ΔN*, *ΔR*, and *M*. Dot position on y-axis indicates fold enrichment, dot size indicates −log10(FDR), and labels highlight significant enrichment in persistent chromatin engram signatures, especially ΔN_28_. The OCT::SOX–AP-1 joint motif was defined as a given distal peak having both OCT::SOX and AP-1 motifs. **d)** TF-encoding genes with concordant evidence from motif enrichment and gene expression. Heatmap shows log2 fold changes for TF genes in *ΔR* and *M* expression estimates, with selected motif families labeled on the left. **e)** Motif cluster enrichment across DEG-linked distal peak sets separated by regulatory quantity and direction. Each point represents a motif cluster, y-axis indicates median enrichment, point size indicates the number of significant motifs in the cluster, and lines connect related motif clusters across regulatory quantities. Motif families enriched in recall response, engram modulation, and no-recall engram chromatin signatures include activity bZIP/AP-1, homeobox, POU/TALE, MEF2/HSF, FOX/SOX, ETS, bHLH/E-box, NFI, NR/ZNF, and T-box/TEAD. **f)** Distal *ΔN_28_*is distinct from the other distal chromatin signatures and is slightly anti-correlated with the early distal engram signature (*ΔN_7_*), whereas gene expression signatures show broader positive coupling across contrasts. Heatmaps show pairwise correlations between decoder weight vectors for *ΔN_7_, M_7_, ΔN_28_*, and *M_28_* within distal chromatin features (left) and within gene expression features (right). Each row and column represents a quantity-specific decoder weight vector, and each cell shows the correlation between the corresponding pair of vectors. In distal chromatin, *ΔN_7_*is strongly aligned with *M_7_* and *M_28_*, and *M_7_* is strongly aligned with *M_28_*, indicating that early engram-associated distal accessibility resembles both early and late modulation-associated distal programs. By contrast, distal *ΔN_28_* separates from this shared axis, showing weak or negative correlation with *ΔN_7_* and *M_7_* and little correlation with *M_28_*. In gene expression, all quantity-specific signatures are positively correlated, with especially strong similarity between *M_7_* and *M_28_*, indicating that transcriptional weight directions are less separated than distal chromatin weight directions. * p < 0.05; ** p < 0.01; *** p < 0.001 for a null of no correlation. **g)** Relationship between distal chromatin *ΔN_28_* decoder weights and gene expression *M_28_* decoder weights without correcting for gene expression *ΔN_28_* decoder weights. Each point represents a gene or linked feature, with selected genes labeled. The scatter compares persistent distal chromatin engram signatures with later transcriptional modulation. Top 5 genes in concordance are highlighted in red and labeled.

**Supplementary Figure 5.**
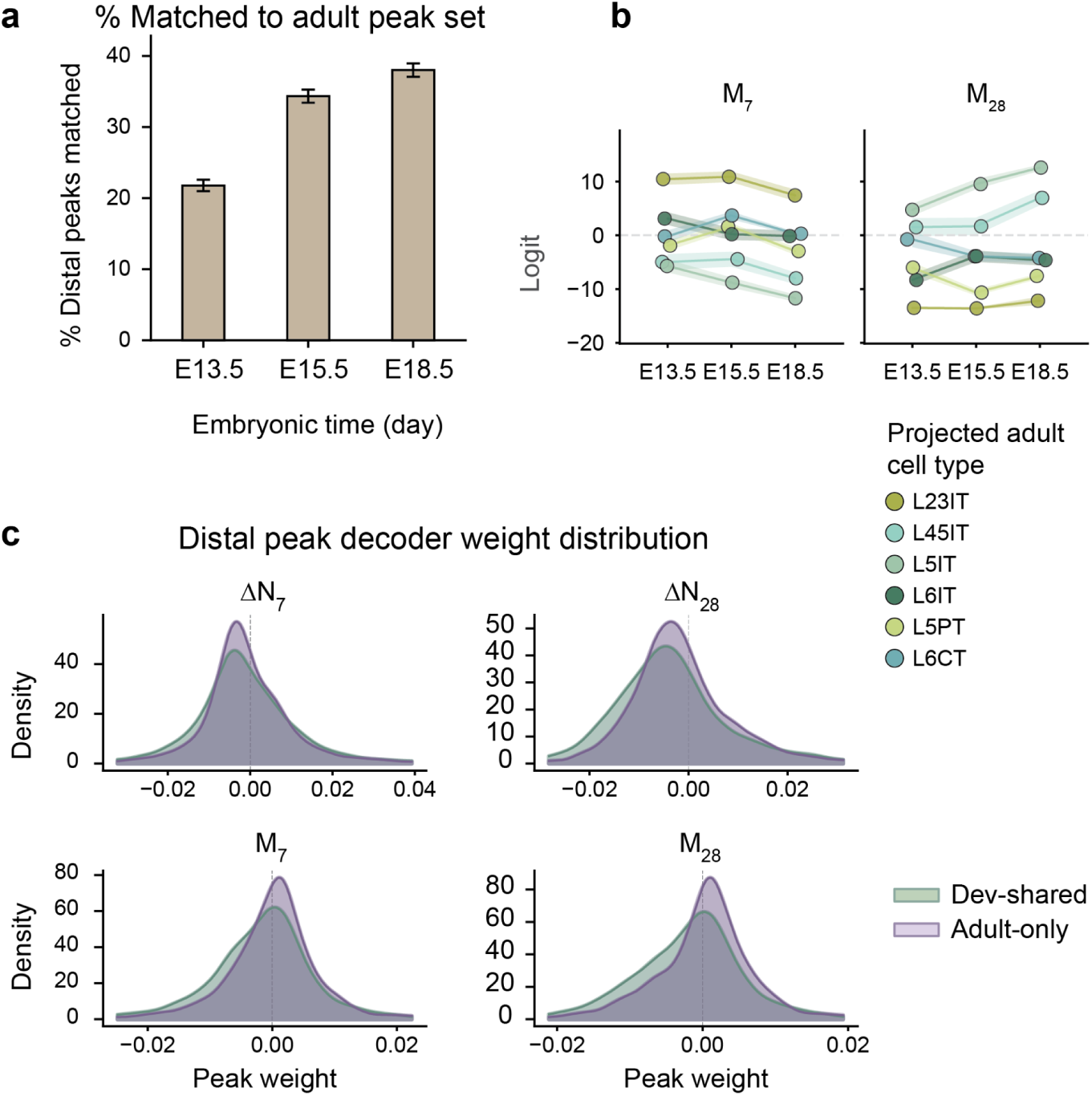
Adult engram and modulation chromatin programs partially reuse embryonic regulatory elements. **a)** Fraction of adult distal peaks matched to embryonic developmental peak sets increases from E13.5, E15.5, to E18.5. Bars show the percentage of adult distal peaks that overlap accessible chromatin regions detected at each embryonic time point, indicating that a subset of adult regulatory elements used in engram and modulation analyses is shared with developmental chromatin programs. Error bars represent 95% Wilson confidence intervals. **b)** Projection of *M_7_* and *M_28_* distal decoder weights onto embryonic developmental time points. Lines connect projected cell types across developmental stages, showing whether peaks associated with engram modulation in adult mPFC align with temporally changing embryonic accessibility programs. Colors represent projected cell types as in Fig. 4b. Shaded ribbons represent standard errors over cross-validation folds. **c)** Distribution of distal peak decoder weights for development-shared and adult-only peaks across *ΔN_7_, ΔN_28_, M_7_*, and *M_28_*. Density plots compare peak weights for distal elements that are shared with embryonic developmental peak sets versus peaks detected only in the adult dataset.

**Supplementary Figure 6.**
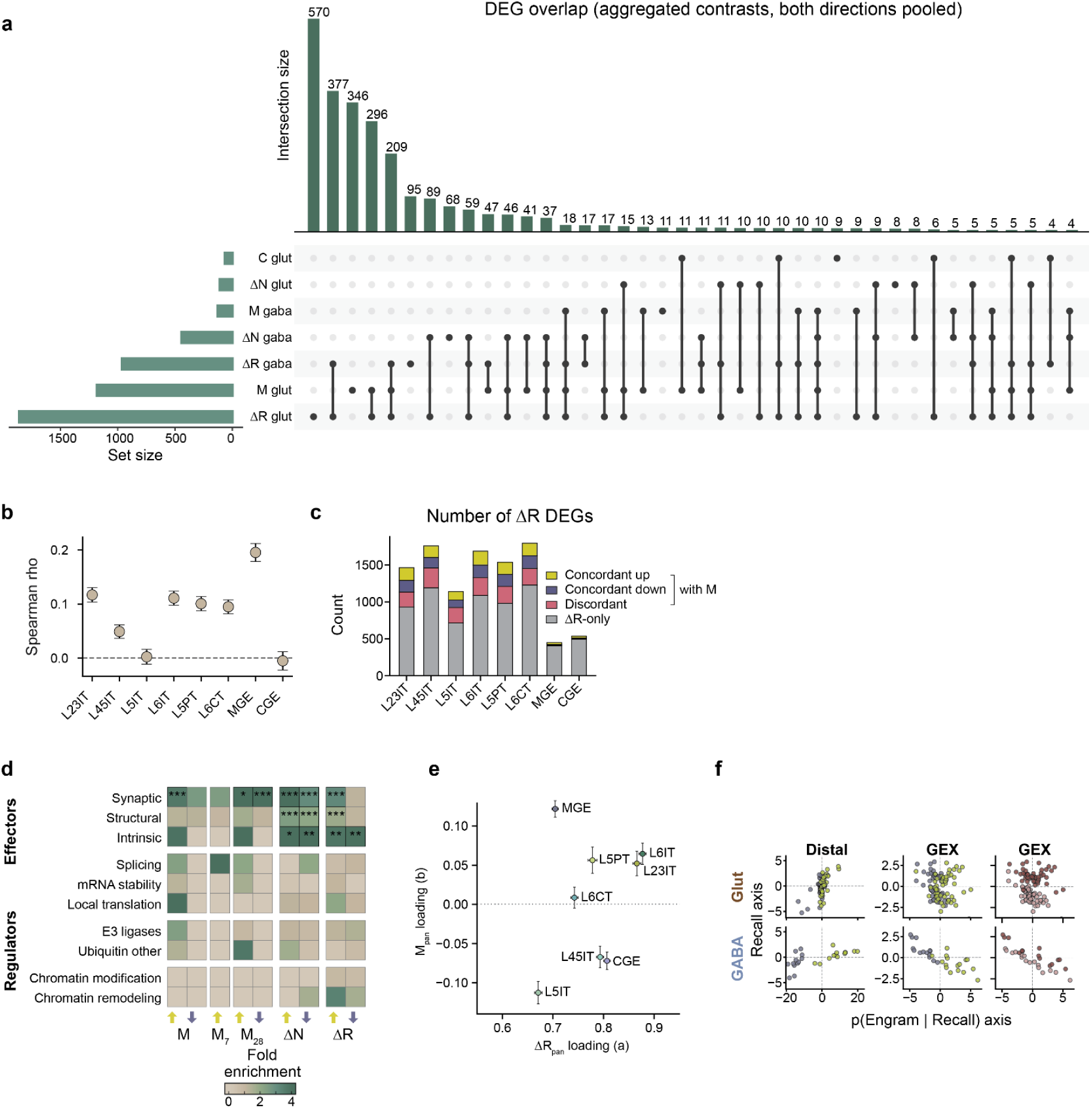
Engram modulation is weaker in GABAergic neurons despite shared recall-response programs. **a)** Overlap of DEGs across glutamatergic and GABAergic regulatory quantities. UpSet plot shows intersections among *ΔR, M, ΔN*, and *C* DEG sets for glutamatergic and GABAergic neurons, with directions pooled across up- and down-regulated genes. Horizontal bars show total set sizes, vertical bars show intersection sizes, and connected dots indicate the DEG sets contributing to each intersection. **b)** Correlation between *ΔR* and *M* gene expression log2 fold changes across matched cell types. Each point represents a neuronal cell type, and the y-axis shows Spearman’s rho between gene expression LFC effect sizes of aggregated *ΔR* and *M*, derived from the DESeq2 model as in Fig. 2. Error bars indicate the standard error of Spearman’s rho, estimated via Monte Carlo simulation (n=1000) by sampling from each gene’s log_2_ fold change distribution using its shrinkage-derived standard error. **c)** Number of recall-response DEGs across neuronal cell types. Stacked bars show genes significant for *ΔR* in each cell type, separated into genes concordant with the pan-cell-class *M* direction, discordant with the *M* direction, or significant only in *ΔR* but not *M*. Glutamatergic cell types were compared to pan-glutamatergic *M*, and GABAergic to pan-GABAergic *M*. **d)** Functional enrichment of DEGs in GABAergic neurons. Heatmap shows fold enrichment for curated effector and regulator categories across *M, M_7_, M_28_, ΔN*, and *ΔR*. Colormap, fold enrichment. * FDR < 0.05; ** FDR < 0.01; *** FDR < 0.001. **e)** Relationship between loadings in pan-cell-class recall-response and engram-modulation for cell-type-specific recall-response LFC vectors. Points represent neuronal cell types, with x-axis showing ΔR_pan_ loading and y-axis showing M_pan_ loading in a linear model of ΔR_cell type_ ∼ ΔR_pan_ + M_pan_. Error bars indicate uncertainty of model estimates. **f)** Projection of glutamatergic and GABAergic cells onto decoder axes. Scatter plots show cells projected onto the logit corresponding to *p(Engram | Recall)* and an orthogonal recall-associated axis for distal chromatin and gene expression features. Panels compare glutamatergic and GABAergic distributions, illustrating that engram-related structure is more separable in GABAergic neurons compared to glutamatergic neurons, and that both cell classes share a common engram-related dimension in the high-dimensional distal chromatin and gene expression space.

**Supplementary Table 1.** Sample metadata.

| Library ID | Condition | Mouse pool ID | Age (week) | Male | Female | Total | Used for DEG analysis | Used for decoder |
| --- | --- | --- | --- | --- | --- | --- | --- | --- |
| YK01 | 7N+ | B2 | 13 | 4 | 0 | 4 | No | Yes |
| YK02 | 7N- | B4 | 13 | 0 | 4 | 4 | Yes | Yes |
| YK03 | 7N+ | B4 | 13 | 0 | 4 | 4 | Yes | Yes |
| YK04 | 7N- | B10 | 20 | 2 | 2 | 4 | Yes | Yes |
| YK05 | 7N+ | B10 | 20 | 2 | 2 | 4 | Yes | Yes |
| YK06 | 7R- | B12 | 15 | 1 | 2 | 3 | Yes | Yes |
| YK07 | 7R+ | B12 | 15 | 1 | 2 | 3 | Yes | Yes |
| YK08 | 7R+ | B14 | 13 | 2 | 2 | 4 | Yes | Yes |
| YK09 | 28N+ | B3 | 16 | 0 | 4 | 4 | Yes | Yes |
| YK10 | 28N- | B7 | 19 | 2 | 2 | 4 | No | Yes |
| YK11 | 28N+ | B7 | 19 | 2 | 2 | 4 | Yes | Yes |
| YK12 | 28N- | B11 | 18 | 2 | 2 | 4 | Yes | Yes |
| YK13 | 28N+ | B11 | 18 | 2 | 2 | 4 | Yes | Yes |
| YK14 | 28R- | B13 | 15 | 2 | 2 | 4 | Yes | Yes |
| YK15 | 28R+ | B13 | 15 | 2 | 2 | 4 | Yes | Yes |
| YK16 | 28R- | B15 | 12 | 2 | 2 | 4 | Yes | Yes |
| YK17 | 28R+ | B15 | 12 | 2 | 2 | 4 | Yes | Yes |

**Supplementary Table 2.** Decoder coefficient matrix.

| | $\Delta R$ | $\Delta N_{\text{mean}}$ | $\Delta N_{\text{time}}$ | $M_{\text{mean}}$ | $M_{\text{time}}$ |
| --- | --- | --- | --- | --- | --- |
| 7N- | 0 | 0 | 0 | 0 | 0 |
| 7R- | 1 | 0 | 0 | 0 | 0 |
| 7N+ | 0 | 1 | -1 | 0 | 0 |
| 7R+ | 1 | 1 | -1 | 1 | -1 |
| 28N- | 0 | 0 | 0 | 0 | 0 |
| 28R- | 1 | 0 | 0 | 0 | 0 |
| 28N+ | 0 | 1 | 1 | 0 | 0 |
| 28R+ | 1 | 1 | 1 | 1 | 1 |

